# The restricted nature of protein glycosylation in the mammalian brain

**DOI:** 10.1101/2020.10.01.322537

**Authors:** Sarah E. Williams, Maxence Noel, Sylvain Lehoux, Murat Cetinbas, Ramnik J. Xavier, Ruslan Sadreyev, Edward M. Scolnick, Jordan W. Smoller, Richard D. Cummings, Robert G. Mealer

**Author notes:** co-senior authors (Richard D. Cummings, Robert G. Mealer). **Corresponding Author:** Robert G. Mealer, M.D., Ph.D., Richard B. Simches Research Center, 185 Cambridge St, 6^th^ Floor, Boston, MA 02114.

## Abstract

Glycosylation is essential to brain development and function, though prior studies have often been limited to a single analytical technique. Using several methodologies, we analyzed Asn-linked (N-glycans) and Ser/Thr/Tyr-linked (O-glycans) protein glycosylation between brain regions and sexes in mice. Brain N-glycans were surprisingly less complex in sequence and variety compared to other tissues, consisting predominantly of high-mannose precursors and fucosylated/bisected structures. Most brain O-glycans were unbranched, sialylated O-GalNAc and O-mannose structures. A consistent pattern was observed between regions, and sex differences were minimal compared to those observed in plasma. Brain glycans correlate with RNA expression of their synthetic enzymes, and analysis of all glycosylation genes in humans showed a global downregulation in the brain compared to other tissues. We hypothesize that the restricted repertoire of protein glycans arises from their tight regulation in the brain. These results provide a roadmap for future studies of glycosylation in neurodevelopment and disease.

## Introduction

Glycosylation regulates nearly all cellular processes and is particularly important in the development and function of the nervous system (Iqbal et al., 2019; Varki, 2017). Glycans have been shown to influence neurite outgrowth (Gouveia et al., 2012), axon guidance (Bonfanti, 2006), synaptogenesis (Parkinson et al., 2013), membrane excitability (Baycin-Hizal et al., 2014; Isaev et al., 2007; Kulkarni et al., 2020; Weiss et al., 2013), and neurotransmission (Kandel et al., 2018; Scott and Panin, 2014) by modulating the structure, stability, localization, and interaction properties of numerous neuronal proteins. Glycans may consist of a single monosaccharide or can be extended into elaborate sugar oligo/polysaccharides (Varki and Kornfeld, 2017). These structures are covalently attached to lipids or certain amino acids of proteins, which designates protein glycans as either N-glycans or O-glycans. Over 300 enzymes work in an elaborate assembly line to generate, attach, and modify these carbohydrate polymers, creating an immense diversity of glycan structures (Rini and Esko, 2017; Varki, 2017).

Despite its complexity, glycosylation is highly regulated; mutations in a single glyco-gene can lead to profound clinical syndromes, collectively termed congenital disorders of glycosylation (CDGs) (Ng and Freeze, 2018). The majority of CDGs present with neurological symptoms including intellectual disability, seizures, and structural abnormalities, illustrating the particular importance of glycosylation in the brain (Freeze et al., 2015). Fine-tuning of the glycosylation pathway can also affect neurophysiology and behavior, illustrated by the association of several glycosylation enzymes to complex human phenotypes such as schizophrenia (Mealer et al., 2020; Schizophrenia Working Group of the Psychiatric Genomics Consortium, 2014), and intelligence (Hill et al., 2018; Joshi et al., 2018).

Prior studies of brain glycosylation have typically focused on a single gene, pathway, epitope, or carrier of interest, providing insight into the roles of specific modifications. Glycolipids have been studied extensively, as they comprise the majority of glycan mass in the brain and are crucial for axon myelination, neuronal survival, and regeneration (Hirabayashi, 2012; Schnaar, 2019; Schnaar et al., 2014). Proteoglycans, composed of a core protein modified by various glycosaminoglycan (GAG) chains, have also been a focus, and are known to be temporally and spatially regulated throughout brain development, serving as guidance cues during cell migration and axon pathfinding (Hussain et al., 2006; Irie et al., 2008; Schwartz and Domowicz, 2018). Less attention has been paid to N- and O-linked protein glycosylation, with a few studies showing the importance of particular modifications such as the Lewis X antigen (Le^X^) (Gouveia et al., 2012; Kudo et al., 1998; Pruszak et al., 2009; Sajdel-Sulkowska, 1998), human natural killer antigen (HNK-1) (Morita et al., 2009; Yamamoto et al., 2002), polysialic acid (Hildebrandt and Dityatev, 2013; Sato and Kitajima, 2019), bisecting GlcNAc (Bhattacharyya et al., 2002; Nakano et al., 2019), and O-mannosylation (Bartels et al., 2016; Stalnaker et al., 2011a, 2011b).

Systematic approaches to capture the diversity of all protein glycans in the brain have been attempted using glycomic analysis (Benktander et al., 2018; Gizaw et al., 2015; Ishii et al., 2007; Ji et al., 2015; Stalnaker et al., 2011a), glycoproteomics (Liu et al., 2017; Riley et al., 2019; Trinidad et al., 2013), microarrays (Zou et al., 2017), western blotting (Simon et al., 2019), and MALDI-Imaging techniques (Powers et al., 2013; Toghi Eshghi et al., 2014). While the majority of these have produced complementary results, they tend to be individually limited by sample size, regional specificity, a single sex, and the technical constraints of a single method. For a more complete picture of brain protein glycosylation, we analyzed the frontal cortex, hippocampus, striatum, and cerebellum of both male and female C57BL/6 mice in the largest such study to date. By comparing results from MALDI-TOF mass spectrometry, tandem mass spectrometry (MS/MS), lectin western blotting, and RNA sequencing, we present a comprehensive portrait of N- and O-glycosylation in the brain, demonstrating a restricted set of glycans and overall downregulation of the pathway.

## Results

### Protein N-glycosylation shows a unique but consistent pattern across the brain

To assess differences in protein glycosylation across brain regions, we first analyzed N-glycans from the cortex, hippocampus, striatum, and cerebellum of 6 adult male wild-type mice using MALDI-TOF mass spectrometry (MS). Representative MALDI spectra from the four regions show a similar pattern of glycans across the brain (**Fig. 1A**), comparable to human cortex and notably distinct from other tissues such as plasma, CSF, and lung (**Fig. S1**). Overall, 95 unique N-glycan masses were annotated across the four regions (**Table S1**). Five of the top 10 most abundant N-glycans in the brain were high-mannose structures, including the most abundant, Man_5_GlcNAc_2_ (Man-5), which comprised nearly half of the total glycan signal (**Fig. 1B**), in contrast to other tissue types which mostly contain more complex, higher molecular weight N-glycans (**Fig. S1**).

**Fig. 1.**
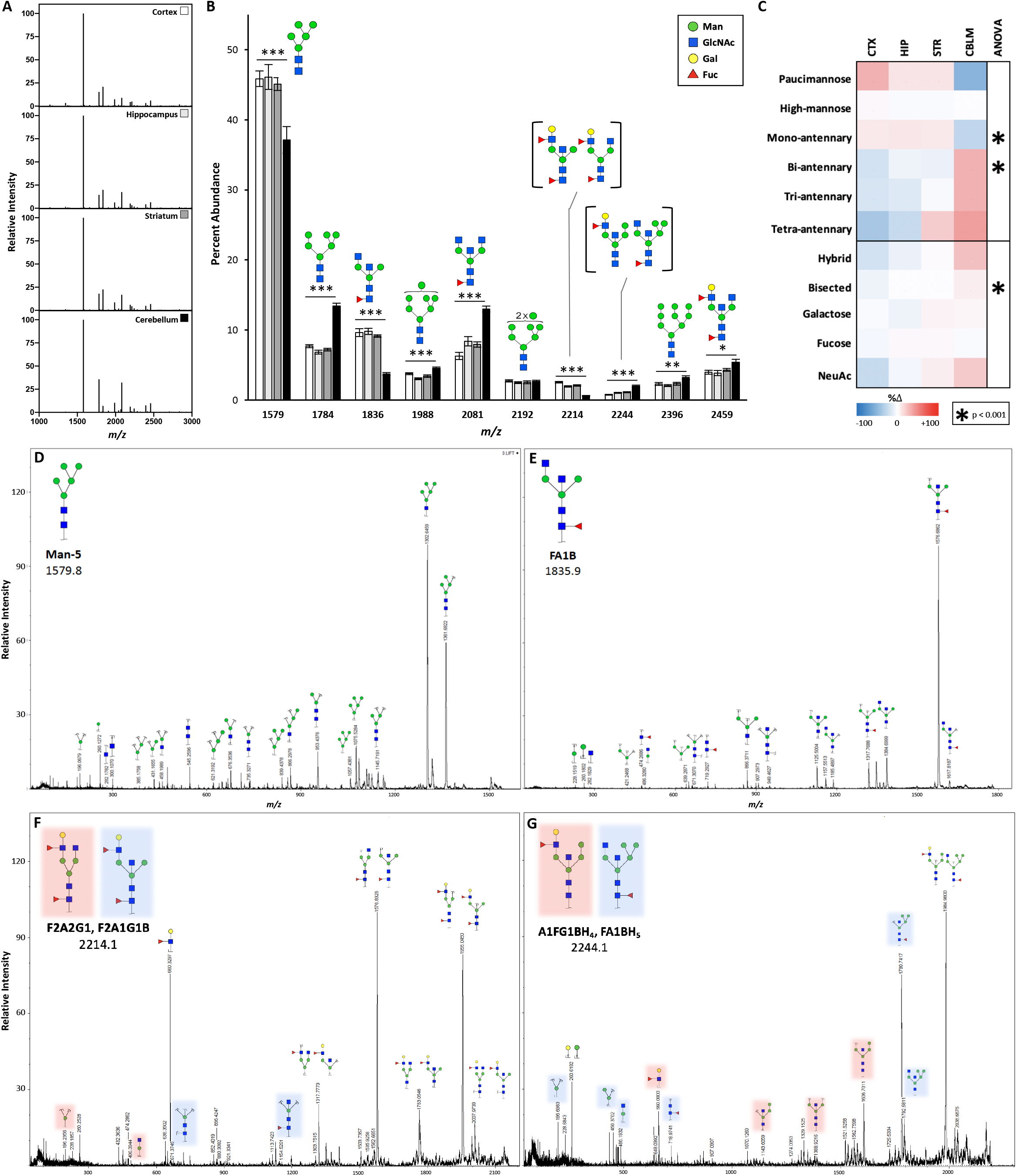
Protein N-glycomics revealed an abundance of high-mannose and fucosylated/bisected structures, confirmed by MS/MS. A) Representative MALDI spectra of protein N-glycans isolated from four brain regions show a similar overall pattern. B) The 10 most abundant N-glycans differ between brain regions (n=6 each). Data presented as mean percent abundance +/- SEM. Corresponding glycan structures are presented above each peak. For ANOVA *df*=(3,20), F_crit_=3.098. C) Categorical analysis of N-glycans demonstrated greater abundance of complex structures in the cerebellum, with a heat map showing percent change from the average of four regions. D) MS/MS analysis of *m/z*: 1579 generated fragment ions all consistent with Man-5. E) Fragmentation of *m/z*: 1836 revealed the presence of a bisected glycan with a terminal mannose. F) Fragmentation of *m/z*: 2214.1 and G) *m/z*: 2244.1 both indicated two distinct glycan structures, consistent with two parent N-glycans at each peak. CTX = cortex, HIP = hippocampus, STR = striatum, CBLM = cerebellum.

For further analysis, individual glycans were categorized by monosaccharide composition or shared structural characteristics such as branching (**Table S2**), and the abundance of these groups were compared between regions. Across the brain, N-glycans were predominantly high-mannose (~60%), fucosylated (~35%), and bisected (~30%) structures (**Table 1**). We noted a low abundance of galactose containing N-glycans (10-15%) and an even smaller amount containing sialic acid (1-3%). Of the few sialylated N-glycans detected, all were modified by the NeuAc form of the sugar.

**Table 1.**
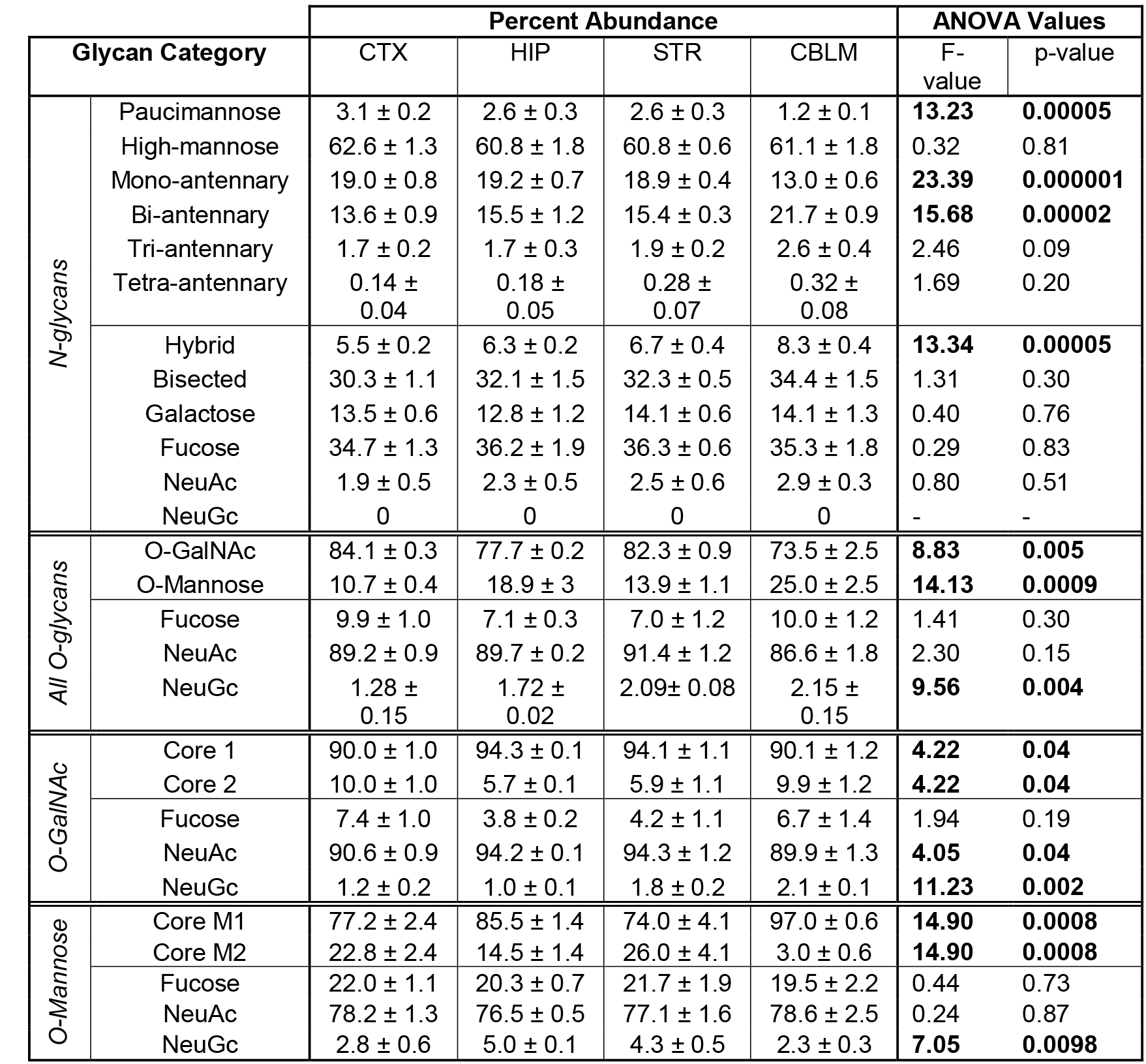
Categorical analysis of N- and O- glycans highlighted regional differences

| Glycan Category |  | Percent Abundance |  |  |  | ANOVA Values |  |
| --- | --- | --- | --- | --- | --- | --- | --- |
|  |  | CTX | HIP | STR | CBLM | F-value | p-value |
| N-glycans | Paucimannose | 3.1 ± 0.2 | 2.6 ± 0.3 | 2.6 ± 0.3 | 1.2 ± 0.1 | **13.23** | **0.00005** |
|  | High-mannose | 62.6 ± 1.3 | 60.8 ± 1.8 | 60.8 ± 0.6 | 61.1 ± 1.8 | 0.32 | 0.81 |
|  | Mono-antennary | 19.0 ± 0.8 | 19.2 ± 0.7 | 18.9 ± 0.4 | 13.0 ± 0.6 | **23.39** | **0.000001** |
|  | Bi-antennary | 13.6 ± 0.9 | 15.5 ± 1.2 | 15.4 ± 0.3 | 21.7 ± 0.9 | **15.68** | **0.00002** |
|  | Tri-antennary | 1.7 ± 0.2 | 1.7 ± 0.3 | 1.9 ± 0.2 | 2.6 ± 0.4 | 2.46 | 0.09 |
|  | Tetra-antennary | 0.14 ± 0.04 | 0.18 ± 0.05 | 0.28 ± 0.07 | 0.32 ± 0.08 | 1.69 | 0.20 |
|  | Hybrid | 5.5 ± 0.2 | 6.3 ± 0.2 | 6.7 ± 0.4 | 8.3 ± 0.4 | **13.34** | **0.00005** |
|  | Bisected | 30.3 ± 1.1 | 32.1 ± 1.5 | 32.3 ± 0.5 | 34.4 ± 1.5 | 1.31 | 0.30 |
|  | Galactose | 13.5 ± 0.6 | 12.8 ± 1.2 | 14.1 ± 0.6 | 14.1 ± 1.3 | 0.40 | 0.76 |
|  | Fucose | 34.7 ± 1.3 | 36.2 ± 1.9 | 36.3 ± 0.6 | 35.3 ± 1.8 | 0.29 | 0.83 |
|  | NeuAc | 1.9 ± 0.5 | 2.3 ± 0.5 | 2.5 ± 0.6 | 2.9 ± 0.3 | 0.80 | 0.51 |
|  | NeuGc | 0 | 0 | 0 | 0 | - | - |
| All O-glycans | O-GalNAc | 84.1 ± 0.3 | 77.7 ± 0.2 | 82.3 ± 0.9 | 73.5 ± 2.5 | **8.83** | **0.005** |
|  | O-Mannose | 10.7 ± 0.4 | 18.9 ± 3 | 13.9 ± 1.1 | 25.0 ± 2.5 | **14.13** | **0.0009** |
|  | Fucose | 9.9 ± 1.0 | 7.1 ± 0.3 | 7.0 ± 1.2 | 10.0 ± 1.2 | 1.41 | 0.30 |
|  | NeuAc | 89.2 ± 0.9 | 89.7 ± 0.2 | 91.4 ± 1.2 | 86.6 ± 1.8 | 2.30 | 0.15 |
|  | NeuGc | 1.28 ± 0.15 | 1.72 ± 0.02 | 2.09 ± 0.08 | 2.15 ± 0.15 | **9.56** | **0.004** |
| O-GalNAc | Core 1 | 90.0 ± 1.0 | 94.3 ± 0.1 | 94.1 ± 1.1 | 90.1 ± 1.2 | **4.22** | **0.04** |
|  | Core 2 | 10.0 ± 1.0 | 5.7 ± 0.1 | 5.9 ± 1.1 | 9.9 ± 1.2 | **4.22** | **0.04** |
|  | Fucose | 7.4 ± 1.0 | 3.8 ± 0.2 | 4.2 ± 1.1 | 6.7 ± 1.4 | 1.94 | 0.19 |
|  | NeuAc | 90.6 ± 0.9 | 94.2 ± 0.1 | 94.3 ± 1.2 | 89.9 ± 1.3 | **4.05** | **0.04** |
|  | NeuGc | 1.2 ± 0.2 | 1.0 ± 0.1 | 1.8 ± 0.2 | 2.1 ± 0.1 | **11.23** | **0.002** |
| O-Mannose | Core M1 | 77.2 ± 2.4 | 85.5 ± 1.4 | 74.0 ± 4.1 | 97.0 ± 0.6 | **14.90** | **0.0008** |
|  | Core M2 | 22.8 ± 2.4 | 14.5 ± 1.4 | 26.0 ± 4.1 | 3.0 ± 0.6 | **14.90** | **0.0008** |
|  | Fucose | 22.0 ± 1.1 | 20.3 ± 0.7 | 21.7 ± 1.9 | 19.5 ± 2.2 | 0.44 | 0.73 |
|  | NeuAc | 78.2 ± 1.3 | 76.5 ± 0.5 | 77.1 ± 1.6 | 78.6 ± 2.5 | 0.24 | 0.87 |
|  | NeuGc | 2.8 ± 0.6 | 5.0 ± 0.1 | 4.3 ± 0.5 | 2.3 ± 0.3 | **7.05** | **0.0098** |

The cerebellum was the most unique of the four brain regions analyzed. Nine of the top ten most abundant N-glycans differed between the cerebellum and other regions, including the most abundant N-glycan, Man-5 (**Fig. 1B**). The cerebellum also displayed significantly less paucimannose and mono-antennary structures, and greater abundance of complex, multi-antennary, and hybrid glycans (**Table 1, Fig. 1C**). Of the other brain regions, the cortex and hippocampus appeared most similar in their composition of N-glycans, and trend towards less complex and branched structures than the cerebellum (**Table 1, Fig. 1C**).

### MS/MS confirms common structural components of N-Glycans

Glycan structures are incredibly diverse, and in some instances, different arrangements of similar monosaccharide components have the same *m/z*, and thus cannot be distinguished by standard mass spectrometry. Therefore, additional analyses are necessary to confirm the structures of glycans detected using MALDI-MS. Tandem mass spectrometry (MS/MS) allows for a single *m/z* peak to be split into fragment ions, which can be analyzed to determine the structure (or structures) of the parent peak in question. We performed MS/MS on several prominent N-glycans from the cortex and cerebellum and obtained complementary results from both regions. We initially presumed several MALDI-MS peaks corresponded to complex, non-bisected N-glycans commonly found in plasma, as reported in prior studies and other tissues (Benktander et al., 2018; Hu et al., 2013; Ji et al., 2015; Jia et al., 2020; Mealer et al., 2019; Mehta et al., 2012; Palmigiano et al., 2016). However, after additional experiments described below, we determined that these peaks primarily represent bisected and/or hybrid N-glycans, further distinguishing the brain N-glycome from that of other tissues.

MS/MS analysis of the most abundant N-glycan, Man-5 (*m/z*: 1579), showed fragment ions consistent with the predicted high-mannose parent structure, including fragments lacking a single mannose (*m/z*: 1361) or a single GlcNAc (*m/z*: 1302), as well as a free hexose (*m/z*: 260) and core GlcNAc (*m/z*: 282, 300) (**Fig. 1D**). The peak at *m/z*: 1835.9, with an abundance of ~10% in most brain regions, could represent several potential structures including a biantennary glycan with two terminal GlcNAc residues and a core fucose (FA2) commonly found in plasma. However, MS/MS demonstrated that this glycan has a bisecting GlcNAc residue and only one antenna (**Fig. 1E**). Several fragment ions from MS/MS results rule in a bisected structure, including a free hexose (*m/z*: 260), which indicates the presence of terminal mannose, and the corresponding structure lacking a terminal mannose (*m/z*: 1617). Additionally, the fragments at *m/z*: 671 and 949 can only result from the fragmentation of a glycan with a bisecting GlcNAc residue. Glycomic analysis of a mouse lacking *Mgat3*, which encodes the only enzyme capable of creating bisected N-glycans (GnT-III), showed a dramatic loss of this glycan in the mutant mouse brain compared to wild-type, providing additional support that *m/z*: 1835 represents a bisected species (Nakano et al., 2019).

Additional MS/MS analyses revealed that some peaks consist of a mixture of at least two unique glycans. For example, MS/MS analysis of *m/z*: 2214 generated fragment ions that indicate the presence of both F2A2G1 and F2A1G1B (**Fig. 1F**). The major peaks *m/z*: 1576 and 1955 could be formed from either parent structure, but the fragment ions at *m/z*: 195 and 486 are specific for F2A2G1, while *m/z*: 671 and 1154 are specific for F2A1G1B. The peak at *m/z*: 2244 contained a mixture of bisected hybrid glycans, with either a Lewis X (Le^X^) epitope and four mannose residues (A1FG1BH4) or a core fucose and five mannose residues (FA1BH5) (**Fig. 1G**). Fragment ions consistent with A1FG1BH4 include the Le^X^ fragment (*m/z*: 660) and the corresponding glycan missing Le^X^ (*m/z*: 1606). Fragment ions consistent with FA1BH5 include fragments missing both the core GlcNAc and core fucose (*m/z*: 1792) as well as several smaller fragments resulting from the tri-mannose hybrid arm (*m/z*: 196, 431 and 450). In sum, our N-glycan MS/MS results are consistent with the predominance of high-mannose, bisected, and fucosylated structures, lower galactose-containing structures, and a very small amount of sialylated N-glycans in the mouse brain.

### Endo H treatment confirms the predominance of high-mannose and hybrid N-glycans in the brain

Given the prevalence of high-mannose N-glycans identified in the brain by MALDI-MS, we sought to confirm this classification with an enzyme that specifically cleaves high-mannose and hybrid structures, known as endoglycosidase H (Endo H). The intensity of individual N-glycans isolated from the cortex using PNGase F (**Fig. 2A**) was compared to those present after Endo H treatment (**Fig. 2B**) and PNGase F digestion following Endo H treatment (**Fig. 2C**). This allowed for the discrimination of structures that are Endo H sensitive, such as high-mannose and hybrid species, and those that are Endo H insensitive, such as paucimannose and complex N-glycans. Of note, PNGase F and Endo H have a different cleavage site on N-glycans, which results in a difference of one GlcNAc residue between the two digestions and prevents the discernment of structures with and without a core fucose following Endo H treatment. Due to this limitation, we primarily focused our comparison on the abundance of PNGase F-released glycans before and after Endo H treatment (**Fig. 2A** vs **2C**) to determine Endo H sensitivity of each parent peak.

**Fig. 2:**
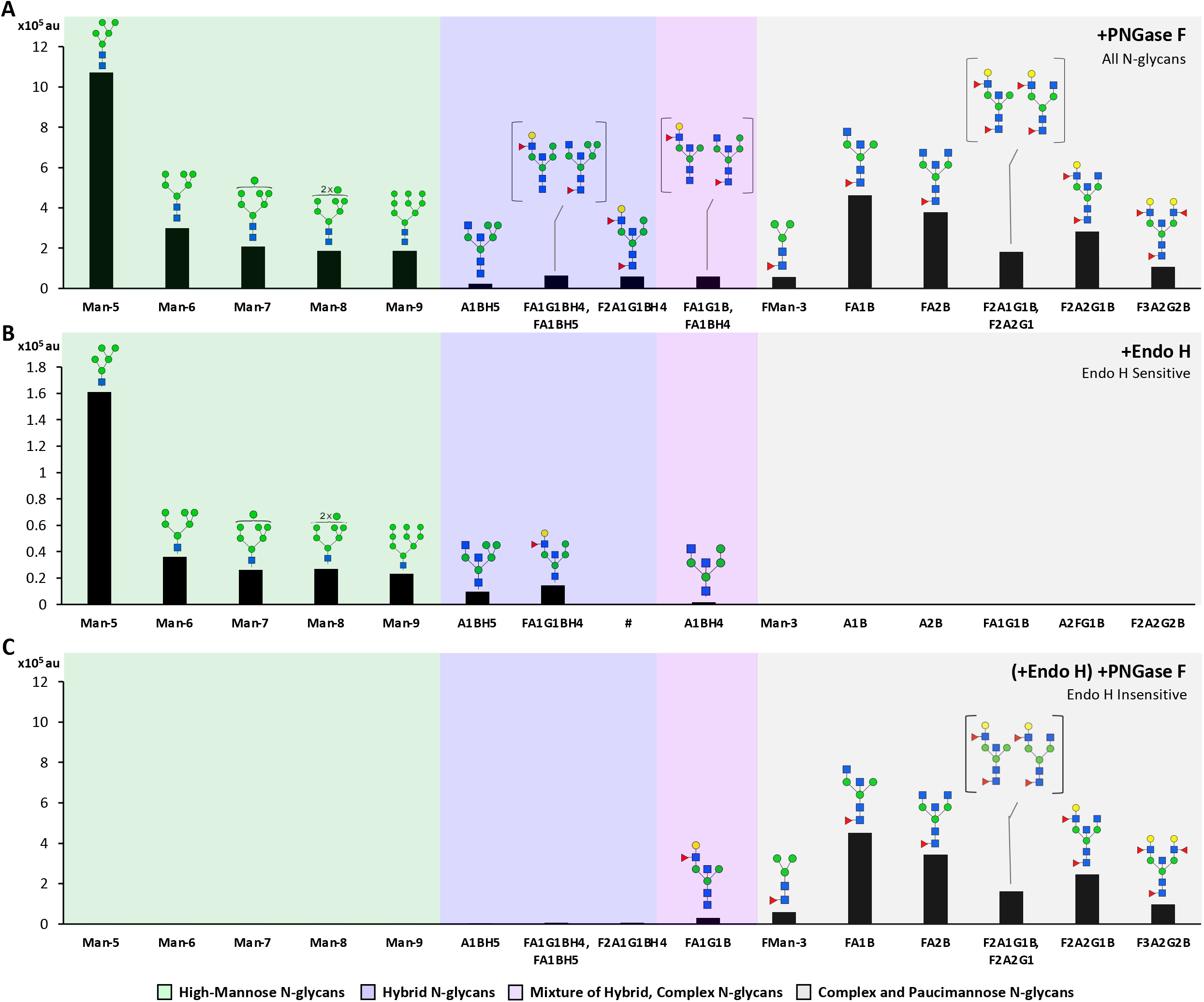
Endo H treatment distinguished high-mannose and hybrid structures from complex and paucimannose N-glycans. The intensity of the 15 most abundant N-glycans in the cortex is shown after treatment with glycosidases, grouped by high-mannose (green), hybrid (blue), mixed (pink), or complex and paucimannose structures (gray). A) Treatment with PNGase F removed all N-glycans. B) Treatment with Endo H removed only high-mannose and hybrid N-glycans. A placeholder (#) is included as the same Endo H fragment can be generated by more than one parent structure if they only differ by the presence of a core fucose. C) PNGase F treatment after Endo H removed complex and paucimannose N-glycans, which were insensitive to Endo H treatment.

Endo H effectively removed 100% of the high-mannose structures present on glycoproteins in the cortex, as none were detected after subsequent PNGase F treatment (**Fig. 2C**). Structures corresponding to Man-5-9 were detected in the Endo H spectra, further supporting this conclusion (**Fig. 2B**). On the contrary, known complex and paucimannose N-glycans were not sensitive to Endo H treatment; these glycans were present at the same relative intensity after the secondary PNGase F treatment (**Fig. 2C**), and no structures corresponding to these glycans were detected in the Endo H spectra (**Fig. 2B**).

Several of the top 15 N-glycan masses identified in the brain have potentially ambiguous structures, as their composition of monosaccharides could form either a hybrid or complex N-glycan structure. For example, the MS peak at *m/z*: 2070 (HexNAc4Hex5) could represent a common plasma N-glycan with two antenna and two terminal galactose residues (A2G2), or a bisected hybrid glycan lacking terminal galactose (A1BH5). Endo H digestion revealed that the N-glycan at *m/z*: 2070 is predominantly the hybrid species A1BH5, as its corresponding mass was detected in the Endo H MALDI spectra (**Fig. 2B**) with minimal signal in the PNGase F spectra after Endo H treatment (**Fig. 2C**). In contrast, another potentially ambiguous glycan (*m/z*: 2214, denoted as F2A2G1, F2A1G1B) was completely insensitive to Endo H digestion, indicating that glycans at this mass do not include a hybrid species, further supported by our MS/MS results (**Fig. 1E**).

In a third unique case, the peak at *m/z*: 2040 was partially Endo H sensitive, indicating a mixture of hybrid and non-hybrid glycans present at this mass. The structure corresponding to the parent hybrid glycan FA1BH4 was detected in the Endo H spectra (A1BH4, **Fig. 2B**) but a small amount of glycan was present in the secondary PNGase F spectra (**Fig. 2C**). MS/MS analysis confirmed the presence of both a hybrid structure and a complex, branched structure present at *m/z*: 2040, which explains why the signal intensity at this mass decreased after Endo H treatment but was not removed entirely (**Fig. S2**).

### Brain O-glycans are primarily sialylated O-GalNAc structures

After removal of N-glycans, we analyzed O-glycans isolated from the remaining brain glycopeptides using MALDI-TOF MS. We identified 26 unique O-glycans in at least one brain region, including both O-GalNAc and O-mannose glycans (**Table S1**). Representative MALDI spectra from the cortex, hippocampus, striatum, and cerebellum showed an overall similar pattern (**Fig. 3A**). However, there are several differences in the abundance of individual O-glycans between brain regions, including the most abundant structure, a di-sialylated core 1 O-GalNAc glycan at *m/z*: 1256 and the most abundant O-mannose glycan, found at *m/z*: 1099 (**Fig. 3B**).

**Fig. 3.**
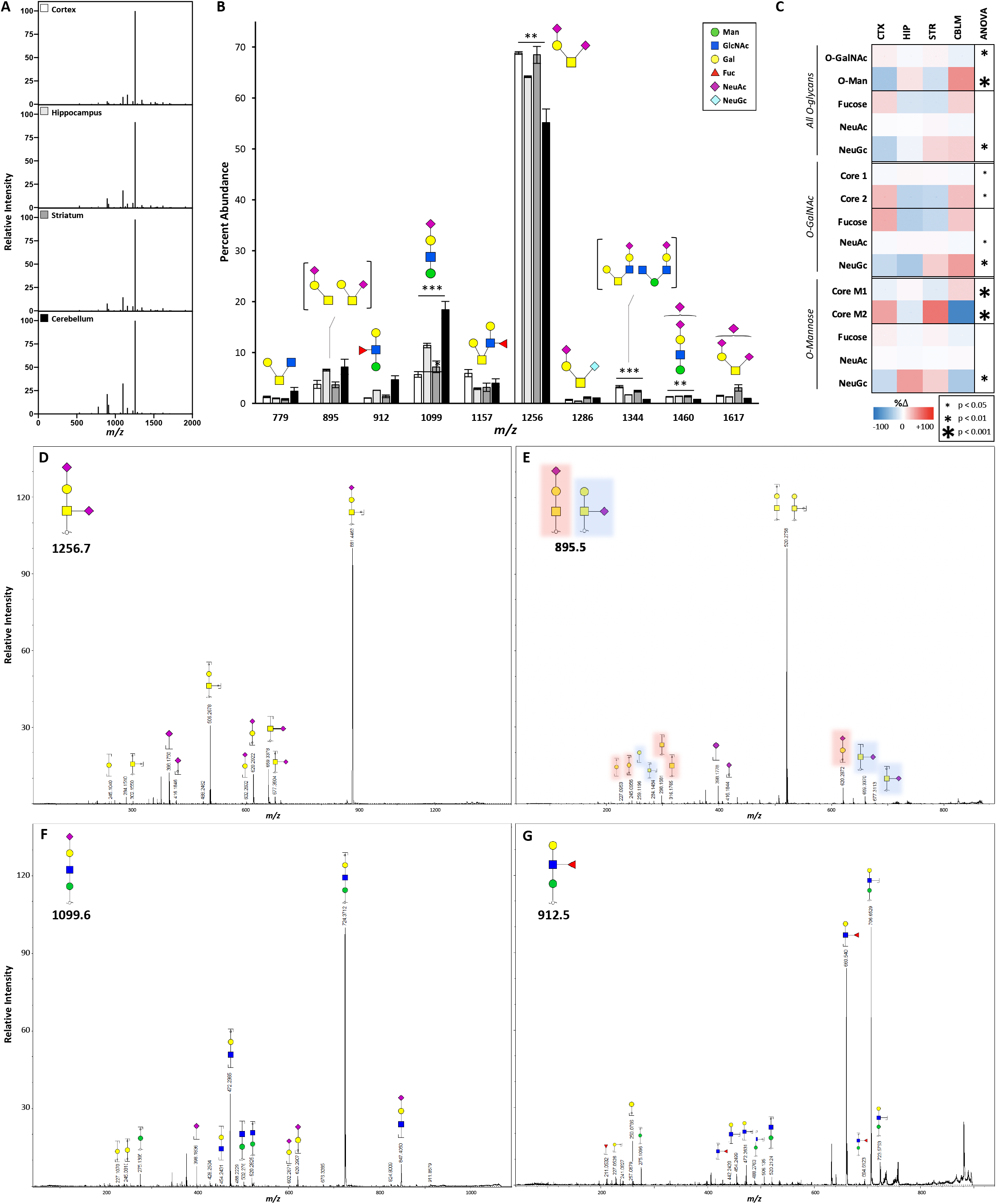
Protein O-glycomics revealed a higher proportion of O-GalNAc glycans compared to O-Man, confirmed by MS/MS. A) Representative MALDI spectra of protein O-glycans isolated from four regions show a consistent pattern across the brain. B) The 10 most abundant O-glycans include both O-GalNAc and O-mannose type structures (CTX=3, HIP=2, STR=4, CBLM=4). Data presented as mean percent abundance +/- SEM. Corresponding glycan structures are presented above each peak. For ANOVA *df*=(3,9), F_crit_=3.862. C) Categorical analysis of O-glycans revealed differences in the abundance of O-GalNAc, O-mannose, and NeuGc-containing glycans between regions. Analyzed independently, O-GalNAc and O-mannose glycans varied in their proportion of core 1, core 2, and sialylated structures. Data presented as a heat map showing percent change from the average of four regions. D) MS/MS analysis of the most abundant O-glycan peak, *m/z*: 1256, generated fragment ions consistent with a di-sialylated core-1 O-GalNAc glycan. D) Fragmentation of *m/z*: 895 identified ions representative of mono-sialylated core-1 O-GalNAc glycans with NeuAc attached to either GalNAc or Gal. E) Fragmentation of *m/z*: 912 and F) *m/z*: 1099 confirm the presence of O-mannose glycans with either terminal sialic acid or fucose.

Analysis of all protein O-glycans stratified by structural components (**Table S3**) revealed that the majority are O-GalNAc-type, comprising 74-84% of the total O-glycan signal across the brain (**Table 1**). The abundance of O-mannose species varies significantly between brain regions, ranging from 11% of all O-glycans in the cortex, to 25% in the cerebellum (**Table 1, Fig. 3C**). In contrast to brain N-glycans, which have a large amount of fucose (~30%) and a paucity of sialic acid (~2%), only ~10% of brain O-glycans are fucosylated, while ~90% are sialylated. Interestingly, we noted very few O-glycans containing both sialic acid and fucose in the brain (<2% in all regions). Simple linear regression of fucosylated vs sialylated O-glycans showed a strong and highly significant negative correlation in both O-GalNAc and O-mannose glycans, suggesting competition between these modifications in the brain (**Fig. S3**). We detected a small amount (1-2%) of O-glycans containing the NeuGc form of sialic acid, consistent with prior studies (Davies and Varki, 2013; Stalnaker et al., 2011a). Among the dominant O-glycans detected, all of the sialylated species contain solely NeuAc (**Fig. 3B**). The most common O-glycan structure, *m/z*: 1256.7, comprises 64% of the total O-glycan abundance and contains two NeuAc residues, while the same structure containing one or two NeuGc residues (*m/z*: 1286 and 1316) was detected at only 0.8% and 0.2% abundance, respectively (**Table S1, S3**).

Of note, not all glycans could be classified as O-GalNAc or O-mannose with confidence, since some peaks corresponded to monosaccharide compositions that could form either type of structure (1-5% of the total glycan signal). For example, *m/z*: 1344, included in the top 10 O-glycans (**Fig. 3B**), could include both O-mannose and O-GalNAc species, as has been reported in a prior study (Stalnaker et al., 2011a). To further analyze brain O-glycans, we took those that were confirmed as O-GalNAc or O-mannose based on MS/MS results (**Fig. 3C-F**) or prior reports (Breloy et al., 2012; Stalnaker et al., 2011a) and normalized the abundance within each O-glycan subtype to sort by structural characteristics (**Table S3** and **Table 1**). Further, we excluded potential structures containing the α-Gal epitope as our results do not confidently rule in its presence, and we did not detect the transcript for its synthetic enzyme α1,3-galactosyltransferase (*Ggta1*) in the brain (Huai et al., 2016).

Analyzed separately, O-GalNAc and O-mannose glycans both varied in their percentage of core 1 and core 2 structures across brain regions (**Table 1, Fig. 3C**). The cortex and cerebellum had higher levels of core 2 O-GalNAc glycans than the hippocampus and striatum, as well as lower amounts of NeuAc-containing glycans, since most of the core 2 O-GalNAc glycans do not contain sialic acid (*m/z*: 1157, 779). Though the cerebellum had overall more O-mannose glycans than the other brain regions, it has nearly ten-fold less core M2 structures, and a corresponding higher proportion of core M1 structures (**Table 1**).

### MS/MS confirms the presence of both O-GalNAc and O-mannose glycans in the brain

Several of the peaks detected from brain O-glycan samples could correspond to either an O-GalNAc or an O-mannose glycan, so we performed MS/MS of highly abundant O-glycans to confirm their structural composition (**Fig. 3D-G**). The most abundant O-glycan peak across all brain regions was *m/z*: 1256, predicted to represent a core 1 O-GalNAc structure modified by two NeuAc residues. We confirmed the structural composition of this glycan, identifying fragments containing NeuAc bound to both the Gal (*m/z*: 620) and GalNAc residues (*m/z*: 659) (**Fig. 3D**). The peak at *m/z*: 895 also contains a core 1 O-GalNAc glycan, modified by one NeuAc residue, and MS/MS results indicated two structural conformations present. The fragment ions at *m/z*: 620, 316, 298, 245, and 227 all indicate the presence of a core 1 O-glycan with NeuAc attached to the Gal residue. Additionally, the fragments at *m/z*: 677, 659, 284, and 259 correspond to a parent glycan with the sialic acid attached to the core GalNAc residue (**Fig. 3E**).

The second most abundant O-glycan in the brain, at *m/z*: 1099, could correspond to three possible parent glycans: an O-GalNAc structure modified with one NeuAc and two Gal residues, an O-GalNAc structure with one Gal, one Fuc, and one NeuGc residue, or an O-mannose glycan extended by GlcNAc, Gal and NeuAc. Fragmentation of *m/z*: 1099 revealed fragment ions exclusive to the O-mannose structure, including core Man (*m/z*: 275) and the corresponding trisaccharide from this cleavage (*m/z*: 847; **Fig. 3F**). Glycomic analysis of a mouse lacking *Pomgnt1*, which encodes the enzyme necessary for extension of O-mannose glycans, revealed a complete loss of *m/z*: 1099 in the mutant mouse brain, supporting the classification of this peak as an O-mannose structure (Stalnaker et al., 2011a).

The peak at *m/z*: 912 was also absent in the *Pomgnt1* knock-out mouse brain, providing evidence that it contains an O-mannose type glycan with a Le^X^ extension, as opposed to a fucosylated O-GalNAc glycan with the same mass. Our MS/MS results confirmed the structure of the O-mannose glycan at *m/z*: 912, with the identification of fragment ions such as the core Man and Le^X^ epitope (*m/z*: 275, 660; **Fig. 3G**). Of note, the *Pomgnt1* knock-out mouse brain retained O-GalNAc glycans, including *m/z*: 1256 and 895, corroborating our determination of these glycans as O-GalNAc-type (Stalnaker et al., 2011a).

### Sex-specific differences in protein glycosylation are minimal in the brain compared to plasma

Previous studies of the brain glycoproteome have primarily focused on mice of a single sex (Ji et al., 2015; Liu et al., 2017; Riley et al., 2019; Toghi Eshghi et al., 2014; Zou et al., 2017). However, it is known that mice show both strain specific and sex specific differences in glycosylation of plasma proteins (Reiding et al., 2016). We used MALDI-TOF glycomics to compare the N-glycome of plasma, cortex, and cerebellum in male and female mice, confirming strong sex differences in the plasma N-glycome but only subtle variation in the brain.

In plasma from male and female mice, we detected 29 N-glycans consisting predominantly of complex, sialylated structures modified by the NeuGc form of sialic acid (**Table S4, S5**), in agreement with previous reports (Reiding et al., 2016). There were striking sex differences in the plasma protein glycomes; notably, the most abundant N-glycan in male mice was A2G2S2 at *m/z*: 2852, while the most abundant N-glycan in female plasma was the fucosylated form of this same species at *m/z*: 3026 (**Fig. 4A**). In addition to this particular glycan, female mice had a 5-fold increase in all fucosylated structures compared to the male plasma glycome (**Table S6**).

**Fig. 4.**
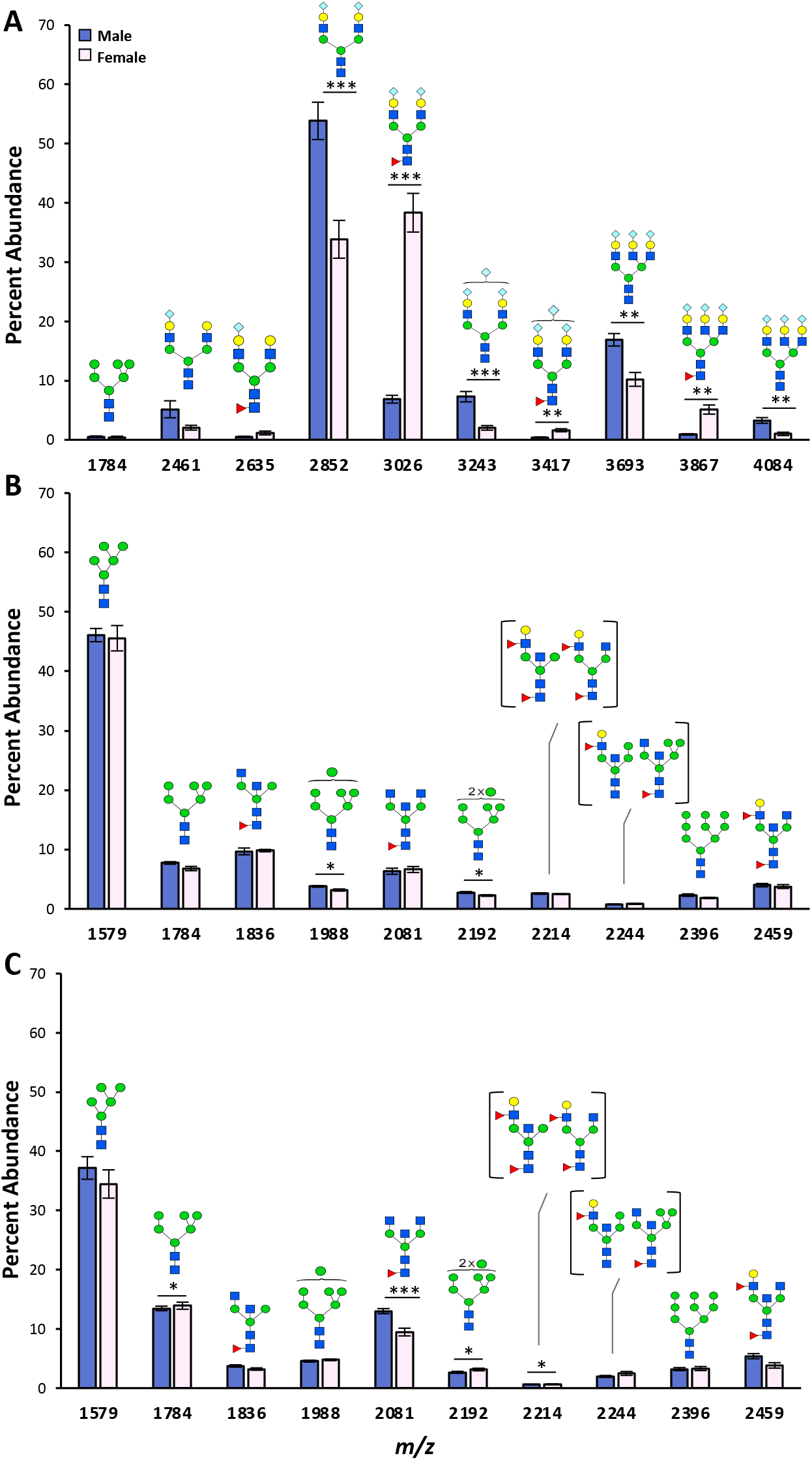
Protein N-glycosylation showed minimal sex differences in the brain compared to plasma. Comparison of the top 10 most abundant N-glycans in plasma (A), cortex (B), and cerebellum (C) reveals differences between sexes, with much greater divergence between male and female mice in the plasma than in either brain region. For plasma samples male=8, female=6. For brain samples male=6, female=4.

In the brain, sex differences in protein N-glycosylation were less pronounced, though significant differences were detected. About 15% of the N-glycans identified in the cortex differed in their abundance compared to the male cortex, including a reduction in Man-7 and Man-8 (**Table S4** and **Fig. 4B**). In the cerebellum, 20% of N-glycans differed from males, including a significant reduction in FA2B at *m/z*: 2081 (**Table S4** and **Fig. 4C**). Based on categorical analysis of these glycans, the female cerebellum showed less biantennary glycans, an increase in sialylation, and an overall trend towards more complex structures compared to the males (**Table S6**). The cortex followed a similar trend but had overall less distinction between sexes. We anticipate that O-glycosylation differences exist between sexes, similar to N-glycosylation. O-glycans from the cortex of two female mice show minor variations in individual glycan abundance compared to the male cortex O-glycans, but these data were not analyzed further due to low sample size (**Fig. S4** and **Table S7**). Though not as pronounced as the differences observed in plasma, these results illustrate that brain protein glycosylation shows some sex-dependence and underscore the importance of analyzing both sexes separately.

### Lectin blotting confirms the high abundance of high-mannose, fucosylated, and bisected N-glycans in the brain

We next performed western blotting of brain glycoproteins using several commercially available biotinylated lectins. Although lectin binding is often not specific for a single epitope, their increased affinity for certain glycan features provides important confirmatory information when complemented with techniques such as glycomics and glycosidase sensitivity. Human plasma was included as a positive control given the abundance of literature on the human plasma N-glycome (Clerc et al., 2016). Glycoproteins were treated with or without PNGase F to determine the relative contribution of N- vs O-glycans to the observed signal. Of note, we detected significant background binding of our fluorescent streptavidin secondary to brain glycoproteins (**Fig. S5**), which likely resulted from high levels of biotin-bound carboxylases in the brain relative to other tissues as previously described (Grant et al., 2019). To reduce this non-specific binding, we pre-cleared the brain lysates by incubation and precipitation with magnetic streptavidin beads, which removed nearly all non-specific binding and allowed for sensitive detection of glycoprotein bands (**Fig. S5**).

Results from lectin blotting agreed with our N-glycomics, indicating high abundances of high-mannose, fucosylated, and bisected structures, with a near absence of galactosylated and sialylated structures (**Fig. 5**). Con A, which binds the core mannose structure of all N-glycans, displayed strong binding in the cortex and cerebellum which was completely sensitive to PNGase F cleavage. GNL, also known as snowdrop lectin, primarily binds extended mannose branches found in high-mannose and hybrid N-glycans. Despite minimal binding in plasma, GNL binding of glycoproteins from both brain regions was robust and PNGase F sensitive, corroborating a predominance of these structures in the brain relative to other N-glycans (**Fig. 1** and **Table 1**). AAL binds fucose in both □(1-3) and □(1-6) linkages of N- and O-glycans. Strong AAL binding was observed in both brain regions and was entirely PNGase F sensitive (**Fig. 5**), suggesting that the bulk of fucose on glycoproteins in the brain was present on N-glycans, in agreement with our glycomics results (**Table 1**). PHA-E, commonly used as a marker for bisected N-glycans, showed strong binding in cortex and cerebellum samples and was PNGase F sensitive. RCA binding, which recognizes galactose in both β(1-3) and β(1-4) linkages, was not detected in brain lysates, but showed strong signal in human plasma, consistent with a relative paucity of galactose terminating glycans in the brain. SNA, also known as elderberry lectin and commonly used to detect glycans with □(2-6)-linked sialic acid, showed only trace binding that was insensitive to PNGase F. This finding is consistent with our glycomics data that only a small minority of N-glycans contain sialic acid (~2%), whereas the majority of O-glycans (>85%) contain at least 1 sialic acid residue (**Table 1**).

**Fig. 5:**
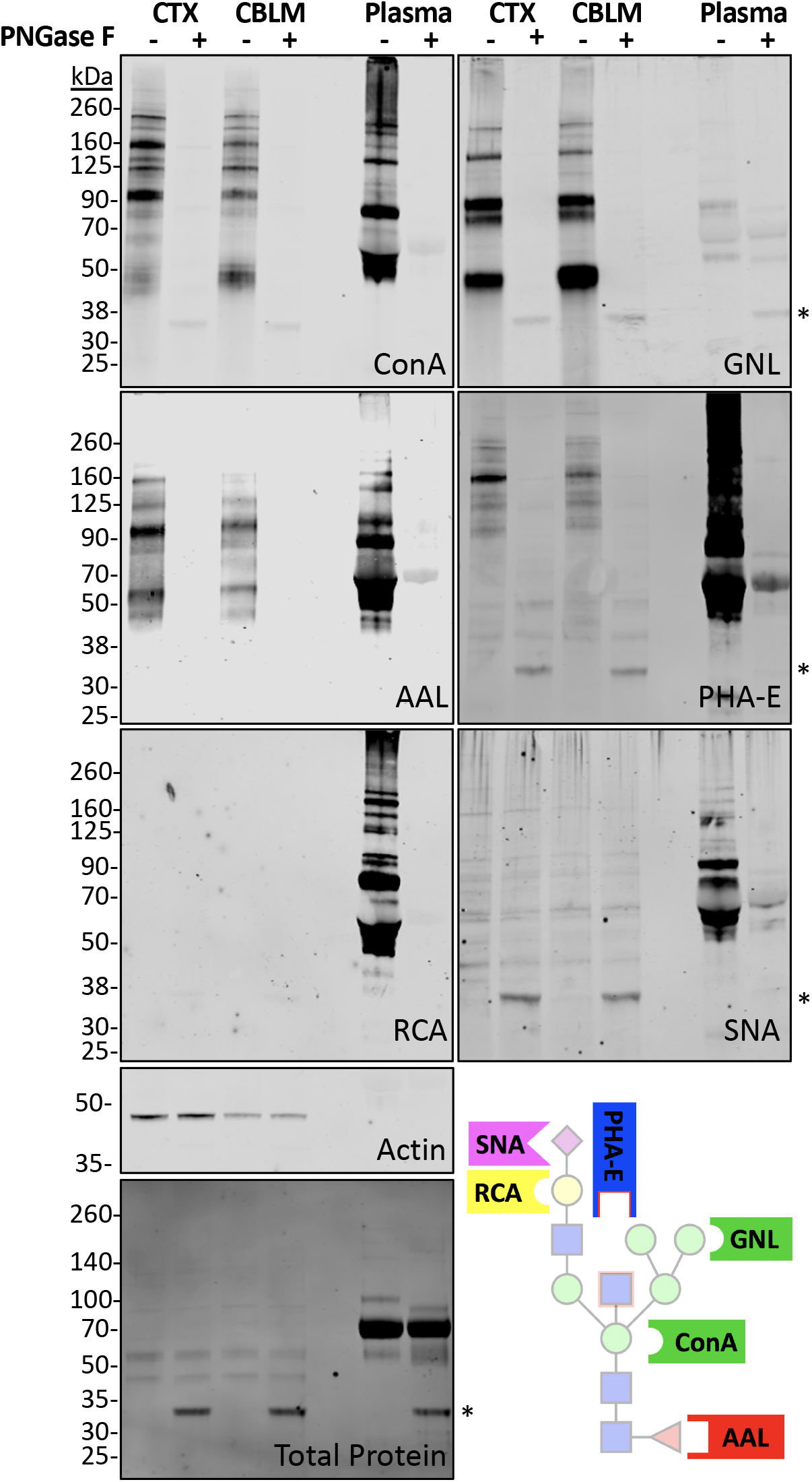
Lectin blotting supported a predominance of high-mannose, fucose, and bisected N-glycans in the brain. Protein lysate from mouse cortex and cerebellum with human plasma as a positive control was treated with or without PNGase F and visualized using biotinylated lectins (Con A, GNL, PHA-E, AAL, RCA, and SNA) in addition to immunoblotting for actin and staining for total protein. Non-specific binding of lectins to PNGase F is noted by an asterisk (*) near 35 kDa. A schematic with common lectin-binding sites is shown for reference.

### Glycosylation gene expression correlates with glycomics and regional differences

We next sought to determine if the expression patterns of glycosylation genes would provide insight into the unique glycome patterns observed in the brain. Comprehensive RNA sequencing and analysis was performed using the contralateral hemispheres of cortex and cerebellum from the same male mice used in our glycomic analysis as previously described (Anders et al., 2015; Dobin et al., 2013; Robinson et al., 2010). We generated a list of 269 known glycosyltransferases, glycosylhydrolases, sulfotransferases, and glycan-related genes based on a previous publication (Joshi et al., 2018) and the Carbohydrate Active Enzymes database (CAZy) (Lombard et al., 2014), after excluding genes whose transcripts were not detected in our experiment (**Table S8**). A comparison between cortex and cerebellum identified 62 differentially expressed glycosylation genes, spanning all synthetic pathways, including protein N-glycans (**Fig. 6A**), O-GalNAc (**Fig. 6B**), and O-Man glycosylation (**Fig. 6C**).

**Fig. 6.**
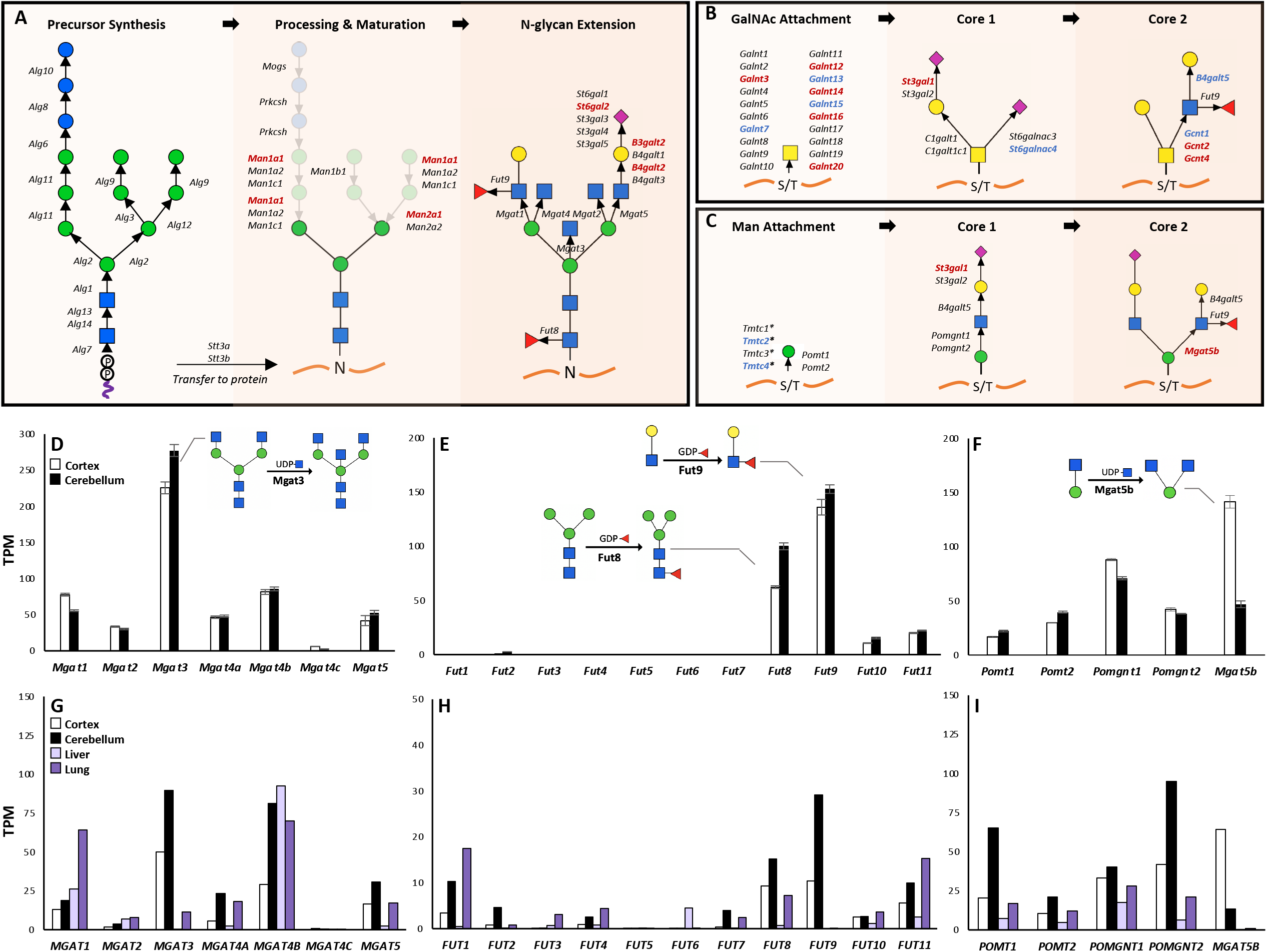
Differential RNA expression between brain regions spans multiple glycosylation pathways and correlates with glycomics results. RNAseq data from cortex and cerebellum (n=4 each) revealed differential expression of enzymes involved in several glycosylation pathways, including the synthesis of N-glycans (A), O-GalNAc glycans (B), and O-mannose glycans (C). Transcripts in red have significantly increased expression in cortex relative to cerebellum, and those in blue are decreased. *\*Tmtc1-4* add O-linked Man but these residues are not extended further. Mouse brain RNAseq results for N-acetylglucosaminyltransferases (D), fucosyltransferases (E), and O-mannose specific enzymes (F) demonstrated high RNA levels of *Mgat3, Fut8, Fut9*, and *Mgat5b*, which correlate with results from glycomics. Human RNA seq data showed a similar expression profile for N-acetylglucosaminyltransferases (G), fucosyltransferases (H), and O-mannose pathway enzymes (I) in the brain between humans and mice, but this pattern is distinct from human liver and lung.

Several correlates between the unique protein glycome and gene expression in the brain were evident. Of the N-acetylglucosaminyltransferases for N-glycans, *Mgat3* levels were much higher than those of branching *Mgat* enzymes (**Fig. 6D**), consistent with the high abundance of bisected N-glycans and the paucity of complex, branched N-glycans. Of the fucosyltransferases, *Fut8* and *Fut9* were most abundant (**Fig. 6E**), correlating with the high amount of core-fucosylated N-glycans and the Le^X^ antigen, respectively. Differential expression of several enzymes between cortex and cerebellum also correlated with the glycomics results. For example, the cortex shows higher expression of *Mgat5b* (**Fig. 6F**), the sole enzyme responsible for synthesis of core-2 O-Man glycans (Haltiwanger et al., 2017), and these structures were several-fold more abundant in this region.

### Human glycosylation genes show a global downregulation in the brain

The unique pattern of protein glycosylation in the mouse brain is mirrored in human samples, which have a similar N-glycan MALDI profile (Fig. S1) and show comparable abundances of high-mannose, bisected, and fucosylated glycans in prior studies (Brown et al., 2019; Gizaw et al., 2016). Gene expression data of human cortex and cerebellum downloaded from the GTEx Portal (Aguet et al., 2019; eGTEx Project, 2017; Lonsdale et al., 2013) revealed several similarities with our RNA expression data from mice for several glycosyltransferase families, including N-acetylglucosaminyltransferases (**Fig. 6G**), fucosyltransferases (**Fig. 6H**), and the enzymes of O-mannosylation (**Fig. 6I**). A comparison to other human tissues with well characterized glycomes, such as liver and lung, illustrated the uniqueness of glycosylation gene expression in the brain. Both brain regions express high levels of *MGAT3* and have a high abundance of bisected N-glycans, while lung and liver have low levels of *MGAT3* and relatively few bisected N-glycans (**Fig. 6G**) (Jia et al., 2020; Mealer et al., 2019; Mehta et al., 2012). The liver and lung have lower levels of nearly all the enzymes for O-Man synthesis and have relatively few of these structures (**Fig. 6I**).

Finally, we compared human glycosylation gene expression in the brain to all other tissues on a global scale (**Fig. 7**). We applied the publicly available GENE2FUNC feature of the FUMA GWAS platform (Watanabe et al., 2017) to a list of 354 glycan-related genes in humans (**Table S9**). Comparison of 54 specific tissue types revealed a distinct pattern of downregulation on the individual gene level across 13 brain regions compared to other tissues (**Fig. 7A, File S1**). Grouped expression analysis of 30 general tissue types showed that the brain is the only region with a significantly down-regulated gene set, and the only region which is significantly different when comparing differences in both directions (**Fig. 7B**). Further analysis of the 13 brain regions as independent tissues shows some regional differences, particularly evident between cortex and cerebellum, though in general the majority of brain regions show an overall downregulation of glycosylation genes (**Fig. S6**).

**Fig. 7.**
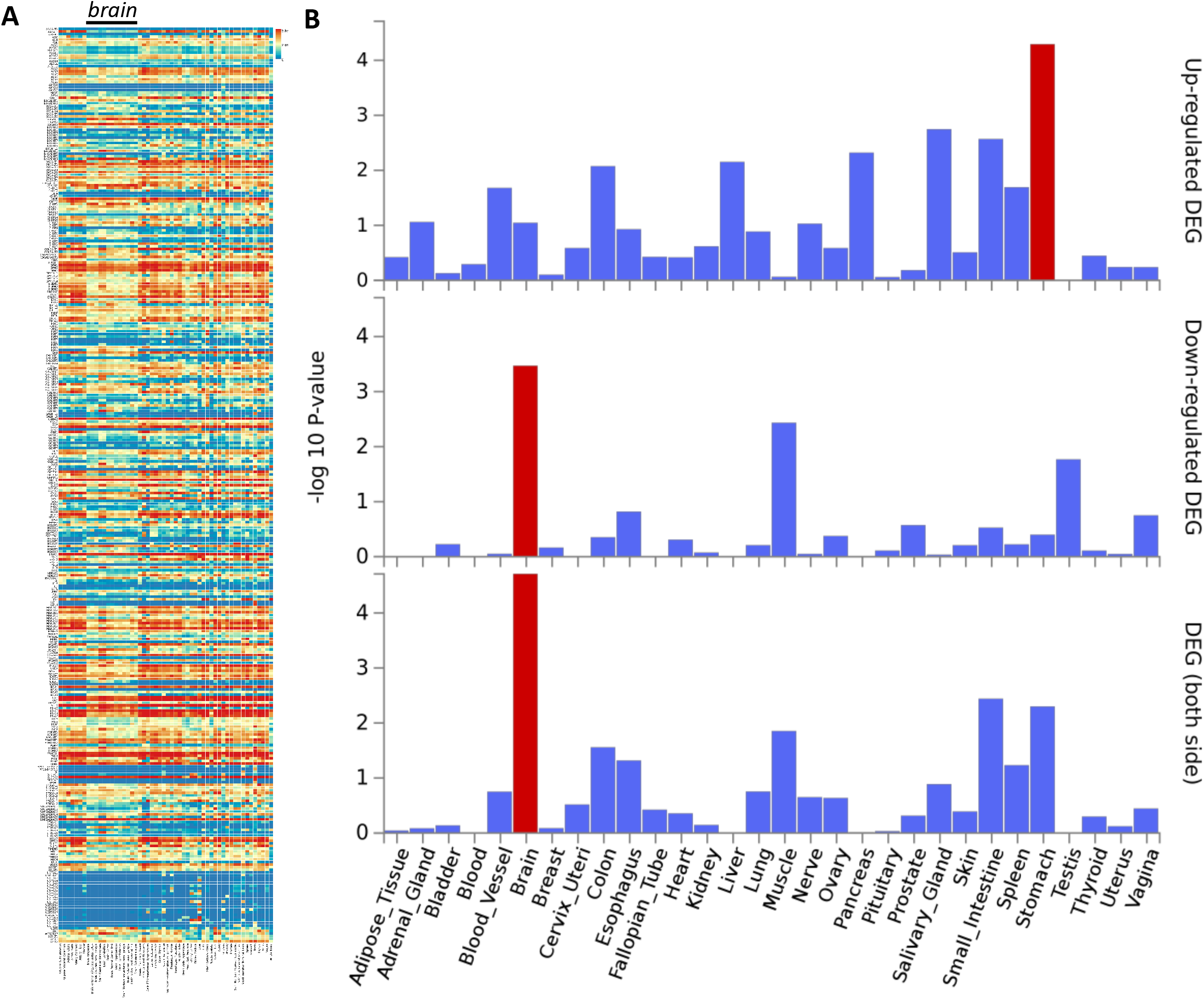
FUMA GENE2FUNC analysis of 354 glycosylation enzymes and related genes in humans revealed a specific downregulation in the brain. A) Heat map demonstrating expression pattern of all glycosylation genes in humans, with a black bar above the 13 columns representing brain regions. Full-size file available in supplement. B) Tissue specific analysis showing down-regulation in the brain compared to all other 29 tissue types, with significantly enriched DEG sets (P_bon_ < 0.05) highlighted in red.

## Discussion

The brain contains millions of cells and billions of connections, creating an unparalleled level of complexity in its development, organization, and regulation. Glycosylation plays a critical role in the establishment and maintenance of this elaborate network, emphasizing the need to understand the unique glycan species involved. Utilizing MALDI-TOF glycomics, MS/MS, lectin blotting, and RNA sequencing, we have generated a comprehensive map of the predominant N- and O-linked protein glycans across multiple brain regions and both sexes of mice. Our findings illustrate a relative simplicity of these structures in the brain and a global downregulation of the pathway, suggesting protein glycan synthesis is tightly controlled.

The overall pattern of brain glycans, in both mouse and human samples, was markedly distinct from those of other tissues. N-glycomics identified predominantly high-mannose and fucosylated/bisected structures in the mouse brain, with few galactosylated, sialylated, or multi-antennary species present, consistent with results from lectin blotting. Both O-GalNAc and O-mannose glycans were detected in the brain, though the former were several-fold more abundant across all brain regions. O-GalNAc and O-mannose glycans consisted primarily of unbranched core 1 structures (as opposed to extended core 2), and in contrast to N-glycans, were almost entirely sialylated. RNA sequencing suggests that gene expression is at least in part responsible for the unique glycome profile observed in the brain. The cortex, hippocampus, striatum, and cerebellum have overall similar glycomes; however, we identified several glycans, glycan classes, and glycosylation enzymes that differ significantly between brain regions, emphasizing the need to study these regions indpendently. The cerebellum was the most unique, with more complex, branched, and hybrid N-glycans, as well as the largest proportion of O-mannose species. We detected relatively few differences in brain protein glycosylation between sexes, in contrast to their distinct plasma N-glycomes, suggesting more conserved regulation of glycosylation in the brain.

The relative simplicity of brain N-glycans is surprising considering their essential physiological roles. High-mannose N-glycans are often considered immature precursor structures but comprise the majority of all N-glycans in the brain. These structures appear to be mature, as they have been detected on the plasma membrane of neurons as well as on extracellular matrix proteins (Kleene and Schachner, 2004; Schmitz et al., 1993; Tucholski et al., 2013a, 2013b). Some studies have demonstrated that these glycans are involved in cell-cell recognition and homeostatic maintenance, governing the interaction properties of NCAM and basigin and influencing neurite and astrocytic outgrowth (Heller et al., 2003; Horstkorte et al., 1993; Kleene and Schachner, 2004). High-mannose N-glycans are also recognized by the mannose receptor (CD206), a microglia specific receptor that can regulate endocytosis and thus may play a role in synaptic pruning (von Ehr et al., 2020; Martinez-Pomares, 2012; Marzolo et al., 1999; Régnier-Vigouroux, 2003).

Of the ~30% of N-glycans in the brain which are not high-mannose structures, the majority (80-90%) are bisected. This may contribute to the lack of extended glycans in the brain, as bisection has been shown to impede subsequent modifications of N-glycans, including galactose and sialic acid, since the additional GlcNAc residue may alter the glycan conformation to prevent interactions with glycosyltransferases (Nagae et al., 2016; Schachter, 1986). *Mgat3* knockout mice develop normally while lacking bisected structures and show a greater relative abundance of complex and modified N-glycans (Nakano et al., 2019). However, high-mannose structures still comprise the majority of N-glycans in the brain of *Mgat3^-/-^* mice, suggesting this molecular brake is only one mechanism in place leading to a low abundance of complex N-glycans.

O-GalNAc glycans can be extensively modified in other organs (Jin et al., 2017; Wheeler et al., 2019) but are limited to mostly sialylated core 1 structures in the brain. Though they comprise the majority of brain O-glycans, the functional roles of O-GalNAc structures are not well understood in the nervous system. A recent case series identified mutations in GALNT2, one of the 20 enzymes capable of attaching the core GalNAc residue to a serine or threonine, as the cause of a novel CDG (Zilmer et al., 2020). Symptoms include intellectual disability, epilepsy, insomnia, and brain MRI abnormalities, and rodent models of *Galnt2* knockout also displayed neurologic abnormalities consistent with a functional role of Galnt2-mediated glycosylation in the brain.

O-mannose structures are better understood in terms of their protein carriers and physiological functions, despite their lower abundance (Darula and Medzihradszky, 2018; Stalnaker et al., 2011a, 2011b). One common carrier is α-dystroglycan, studied extensively in congenital muscular dystrophies, though knockout studies have shown that there are many other proteins modified by O-mannose in the brain (Endo, 2015; Stalnaker et al., 2011a). Extended O-mannose glycans, including those harboring the HNK-1 and Le^X^ epitopes, have been identified on components of perineuronal nets, extracellular matrix structures involved in cell adhesion and neurite outgrowth (Morise et al., 2014; Pacharra et al., 2013; Yaji et al., 2015). Core M2 glycans have only been reported in the brain, where the key synthetic enzyme MGAT5B is highly enriched, and regulate remyelination, astrocyte activation, and oligodendrocyte differentiation (Inamori et al., 2003; Kanekiyo et al., 2013; Kaneko et al., 2003; Kizuka et al., 2014; Lee et al., 2012). A unique mono-O-mannose glycan on members of the cadherin family has been recently described, and is necessary for the cell-adhesion function of these proteins (Lommel et al., 2013; Vester-Christensen et al., 2013). This O-Man attachment is catalyzed by a novel family of O-mannosyltransferases known as TMTC1-4, rather than the canonical POMT-initiated O-mannose pathway, and is not extended further than the core Man residue (Larsen et al., 2017, 2019).

Of note, we do not identify monosaccharide modifications, including mono-O-Man, mono-O-Fuc, or mono-O-GlcNAc, despite brain expression of their synthetic enzymes (*Tmtc1-4, Pofut1-2*, and *Ogt*). Such modifications may be present at a lower abundance relative to extended O-GalNAc and O-mannose glycans in the brain, as previous studies have primarily used enrichment strategies for their isolation (Holdener and Haltiwanger, 2019; Larsen et al., 2017; Thompson et al., 2018).

While less than 3% of brain N-glycans are modified by sialic acid, almost all of the O-glycans detected in this study are sialylated. This charged carbohydrate residue has been studied extensively in the context of brain development and disease (Sato and Kitajima, 2019). Sialic acid has been investigated as a regulator of phagocytosis, as microglia express several siglec-type receptors that recognize sialic acid and trigger an inhibitory response in the cell upon binding (Linnartz and Neumann, 2013; Siddiqui et al., 2019). Enzymatic removal of sialic acid from neurons in culture decreases siglec binding, increases engulfment by microglia, and potentiates complement deposition, a key regulatory step in microglial-mediated synaptic pruning (Linnartz et al., 2012; Schafer et al., 2012; Schizophrenia Working Group of the Psychiatric Genomics Consortium et al., 2016; Stevens et al., 2007; Wielgat and Braszko, 2012). Another carrier of sialic acid in the brain is PSA-NCAM, which can harbor up to 400 sialic acid residues and is critical in brain development and neuronal migration (Nakata and Troy, 2005; Schnaar et al., 2014). We did not identify this structure in our samples likely due to its large size and low abundance in the adult brain (Nacher et al., 2013; Quartu et al., 2008; Seki and Arai, 1993).

In sum, we present a comprehensive picture of protein N- and O-glycosylation in the mouse brain. Our results highlight unique glycan compositions and distinct regulatory mechanisms across several brain regions, tissue types, and sexes in the largest sample size to date. We emphasized the most abundant N- and O-glycans in the brain and their potential physiological roles, but this makes no assumption of the function or importance of structures that exist at very low abundance. Subtle changes in glycosylation can lead to major consequences at the protein, cell, and circuit level, so it is essential to understand how such variation is regulated at the genetic (Joshi et al., 2018), epigenetic (Kizuka et al., 2016), transcriptional (Neelamegham and Mahal, 2016), developmental (Ishii et al., 2007; Simon et al., 2019), regional (Gizaw et al., 2015; Powers et al., 2014; Toghi Eshghi et al., 2014), and organismal levels (Brown et al., 2019; Gizaw et al., 2016; Yamakawa et al., 2018). The contribution of glycosylation to health and disease has been appreciated in many contexts, especially the nervous system (Reily et al., 2019). In addition to neurologic symptoms of CDGs (Freeze et al., 2015), complex neuropsychiatric phenotypes are linked to glycosylation (Hill et al., 2018; Joshi et al., 2018; Williams et al., 2020). For example, several glycosyltransferases and a missense variant in *SLC39A8* are associated with schizophrenia, emphasizing the need for a more detailed understanding of protein glycosylation as it relates to development and disease in the brain (Mealer et al., 2020).

## Author contributions

SEW performed glycomics experiments, statistical analysis, assisted in RNA analysis, and wrote the manuscript with RGM

MN performed lectin blotting, assisted in RNA analysis, and wrote portions of the manuscript

SL assisted with experimental methods and advised on analysis of glycans

MC performed analysis of RNAseq data

RX provided mouse samples for analysis and assisted with project design

RS oversaw RNAseq analysis and wrote related portions of the manuscript

EMS initiated the project and coordinated collaborations

JWS co-supervised SEW and helped conceptualize the project

RDC co-supervised SEW and MN, oversaw all experimental analyses, and helped conceptualize the project

RGM co-supervised SEW and MN, performed glycomics experiments, lectin blotting, RNA purification and analysis, statistical analysis, helped conceptualize the project, and wrote the manuscript

All authors contributed feedback and edits to the manuscript

## Acknowledgements

This work was supported by a foundation grant from the Stanley Center for Psychiatric Research at the Broad Institute of Harvard/MIT (awarded to RGM) and NIH grant P41GM103694 (awarded to RDC).

## STAR Methods

### RESOURCE AVAILABILITY

#### Lead Contact

Further information and requests should be directed to the Lead Contact, Robert Mealer.

#### Materials Availability

This study did not generate any new reagents.

#### Data and Code Availability

This study did not generate any code, and datasets generated from the current study are available from the lead contact upon request.

### EXPERIMENTAL MODEL AND SUBJECT DETAILS

Fresh post-mortem mouse brain samples were harvested from wild-type mice on a C57BL/6J background originally from The Jackson Laboratory (Cat#000664) after euthanasia with CO_2_. Mice from both sexes were used in this study and were 12 weeks old at the time of tissue harvest, sample size specified for each experiment. No live animals were used in this study. All mice were housed and maintained in accordance with the guidelines established by the Animal Care and Use Committee at Massachusetts General Hospital under protocol #2003N000158. Human Brain Cerebral Cortex Whole Tissue Lysate was purchased from Novus Biologicals (#NB820-59182).

### METHOD DETAILS

#### Sample Preparation

##### Brain dissection

Following euthanasia with CO_2_, whole mouse brain was removed and placed on a clean ice cold plastic surface and rinsed with PBS at 4°C. Four brain regions (frontal cortex, hippocampus, striatum, cerebellum) were isolated from each hemisphere using blunt dissection and placed in 1.5 mL conical tubes, snap frozen in liquid N_2_, and stored at −80°C until further use.

##### Tissue lysis

Frozen brain tissue was lysed in 500μL ice cold lysis buffer (50 mM TRIS, 150 mM NaCl, 1.0% w/v Triton-X, pH 7.6) with protease inhibitor (Roche #46931320019) and dissociated using a hand-held motorized pestle (Kimble #749540), followed by 2 brief pulses of sonication for 10 seconds with a microtip (Qsonica Q700). An additional 500μL of lysis buffer was added to bring the volume to 1 mL, and protein concentration was analyzed using the Pierce BCA Protein Assay Kit (ThermoFisher Scientific #23255).

#### Glycomic Analysis

##### Isolation and purification of glycoproteins

Brain glycoproteins were purified according to standard protocols readily available through the National Center for Functional Glycomics website (www.ncfg.hms.harvard.edu). All buffers were made fresh daily. In brief, 2 mg of protein lysate per sample was dialyzed in 3.5 L of 50 mM ammonium bicarbonate 3 times at 4°C over 24 hours using snakeskin dialysis tubing with a molecular cut-off between 1 and 5 kDa (ThermoFisher Scientific #68035). Samples were lyophilized and then resuspended in 1 mL of 2 mg/mL 1,4-dithiothreitol (DTT) dissolved in 0.6 M TRIS (pH 8.5) and incubated at 50°C for 1.5 hours, followed by addition of 1 mL of 12 mg/mL iodoacetamide in 0.6 M Tris (pH 8.5) and incubated at room-temperature for 90 minutes in the dark. Samples were again dialyzed as described above, lyophilized, and resuspended in 1 mL of 500 μg/ml TPCK-treated trypsin in 50 mM ammonium bicarbonate and incubated overnight (12-16 hours) at 37°C. Trypsin digestion was stopped by addition of ~2 drops 5% acetic acid, and samples were added to a C18 Sep-Pak (200 mg) column (Waters, #WAT054945) preconditioned with one column volume each of methanol, 5% acetic acid, 1-propanol, and 5% acetic acid. Reaction tubes were washed with 1 mL 5% acetic acid and added to the column, followed by an additional 5 mL wash of 5% acetic acid. Each column was placed in a 15 mL glass tube, and glycopeptides were eluted using 2 mL of 20% 1-propanol, 2 mL of 40% 1-propanol, and 2 mL of 100% 1-propanol. The eluted fractions were pooled and placed in a speed vacuum to remove excess organic solvent, followed by lyophilization.

##### Release and purification of protein N-glycans

Lyophilized glycopeptides were resuspended in 200 μL of 50 mM ammonium bicarbonate and incubated with 3 μL of either PNGase F (New England Biolabs, #P0704) or Endo H (New England Biolabs, #P0702S) at 37°C for 4 hours, then overnight (12-16 hours) with an additional 5 μL of enzyme at 37°C. C18 Sep-Pak columns (200 mg) were preconditioned with one column volume of methanol, 5% acetic acid, 1-propanol, and 5% acetic acid and placed in 15 mL glass tubes. Digested samples were loaded onto preconditioned columns, collecting all flow-through, and N-glycans were eluted with 6mL of 5% acetic acid. Eluted fractions were pooled and placed in a speed vacuum to remove the organic solvents and lyophilized overnight. Glycopeptides remaining on the C18 columns were eluted using 2 mL of 20% 1-propanol, 2 mL of 40% 1-propanol, and 2 mL of 100% 1-propanol, placed in a speed vacuum to remove the organic solvents and lyophilized for O-glycan processing.

##### β-elimination and purification of O-glycans

After removing N-glycans from glycopeptides, O-linked glycans were removed using a *β*-elimination reaction according to the standard protocols available through the National Center for Functional Glycomics (www.ncfg.hms.harvard.edu). In brief, lyophilized N-glycan-free glycopeptides were resuspended in 400 μL of 55 mg/mL NaBH4 in 0.1 M NaOH solution and incubated overnight (12-16 hours) at 45°C. β-elimination reaction was terminated by adding 4-6 drops of glacial acetic acid to each sample. Desalting columns were prepared using Dowex 50W X8 ion exchange resin with mesh size of 200-400 (Sigma-Aldrich, #44519) in small glass Pasteur pipettes and washed with 10 mL of 5% acetic acid. Acetic acid-neutralized samples were loaded onto columns, collecting flow through in 15 mL glass tubes. Columns were washed with an additional 3 mL of 5% acetic acid and all flow-through was pooled, placed in a speed vacuum to remove organic solvent and lyophilized. Dried samples were resuspended in 1 mL of 1:9 acetic acid:methanol solution (v/v=10%) and dried under a stream of nitrogen, repeating this step an additional three times. C18 Sep-Pak columns (200 mg) were preconditioned with one column volume of methanol, 5% acetic acid, 1-propanol and 5% acetic acid and placed in 15mL glass tubes. The dried samples were resuspended in 200 μL of 50% methanol and loaded onto the conditioned C18 columns, collecting flow-through. Columns were washed with 4mL of 5% acetic acid and all flow-through pooled, placed in a speed vacuum to remove the organic solvents and lyophilized.

##### Preparation and isolation of plasma N-glycans

Blood samples were collected following CO2 euthanasia and decapitation in a microtainer tube (BD, #365967), and plasma was separated by centrifugation and stored at −80°C until use. Plasma N-glycan profiling was performed as described previously (Mealer et al., 2019). In brief, 5μL of mouse plasma was lyophilized, resuspended in 20 μL 1X Rapid PNGase F buffer (NEB #P0710S), and denatured at 70°C for 15 min. After cooling to room temperature, 1 μL of Rapid PNGase F was added, and incubated at 50°C for 60 min. C18 Sep-Pak columns (50 mg, Waters, #WAT054955) were preconditioned with one column volume of methanol, 5% acetic acid, 1-propanol, and 5% acetic acid and placed in 1.5 mL tubes. PNGase F-treated samples were resuspended in 100 μL of 5% acetic acid and added to the preconditioned columns, collecting all flow-through. The reaction tube was washed with an additional 100 μL of 5% acetic acid which was added to the column, followed by 1 mL of 5% acetic acid, and the entire flow-through was placed in a speed vacuum to remove the organic solvents and lyophilized prior to permethylation as described below.

##### Permethylation of N- and O-glycans

Using a clean, dry mortar and pestle, 21 pellets of NaOH were ground and dissolved into 12 glass pipettes volumes (~3 ml) of DMSO. A fresh slurry of NaOH/DMSO was made daily. 1 mL of the slurry was added to the lyophilized N- and O-glycans in addition to 500 μL of iodomethane (Sigma Aldrich, #289566). Samples were tightly capped and placed on a vortex shaker for 30 minutes at room temperature. After the mixture became white, semi-solid, and chalky, 1 mL ddH_2_O was added to stop the reaction and dissolve the sample. 1 mL of chloroform and an additional 3 mL ddH_2_O were added for chloroform extraction and vortexed followed by brief centrifugation. The aqueous phase was discarded, and the chloroform fraction was washed three additional times with 3 mL ddH_2_O. Chloroform was then evaporated in a speed vacuum. Permethylated glycans were resuspended in 200 μL of 50% methanol and added to a C18 Sep-Pak (200 mg) column preconditioned with one column volume each of methanol, ddH_2_O, acetonitrile, and ddH_2_O. The reaction tubes were washed with 1 mL 15% acetonitrile and added to the column, followed by an additional 2 mL wash of 15% acetonitrile. Columns were placed into 15 mL glass round-top tubes, and permethylated glycans were eluted with 3 mL 50% acetonitrile. The eluted fraction was placed in a speed vacuum to remove the acetonitrile and lyophilized overnight.

##### MALDI-TOF Acquisition

Permethylated glycans were resuspended in 25 μL of 75% methanol and spotted in a 1:1 ratio with DHB matrix on an MTP 384 polished steel target plate (Bruker Daltonics #8280781) as previously described (Mealer et al., 2019). MALDI-TOF MS data was acquired from a Bruker Ultraflex II instrument using FlexControl Software in the reflective positive mode. For N-glycans, a mass/charge (*m/z*) range of 1,000-5,000 kD was collected, and for O-glycans, a range of 500-3,000 kD. Twenty independent captures (representing 1,000 shots each) were obtained from each sample and averaged to create the final combined spectra file. Data was exported in .msd format using FlexAnalysis Software for subsequent annotation. Tandem MS (MS/MS) data was collected using the same instrument for both N- and O-glycans, using the LIFT positive mode, and a +/- 1 Da range from the predicted parent *m/z*, and again represent the sum of twenty independent captures.

##### N- and O-Glycan Analysis

Glycans of known structure corresponding to the correct isotopic mass which had a signal to noise ratio greater than 6 (S/N) in at least one brain region averaged over the grouped samples were annotated using mMass software (Strohalm et al., 2010). This resulted in 95 brain N-glycans, 26 brain O-glycans, and 29 plasma N-glycans. The relative abundance of each glycan was calculated as the signal intensity for each isotopic peak divided by the summed signal intensity for all measured glycans within a spectrum. Although using the isotopic mass for quantification may underestimate the relative abundance of larger glycans given the increased incorporation of Carbon-13, the majority of N- and all of O-glycans in the brain are best represented by the isotopic peak (*m/z* < 2040). Structural assignment of glycans was based on MS/MS results, enzyme sensitivity (PNGase F, Endo H), previously confirmed structures (Breloy et al., 2012; Nakano et al., 2019; Stalnaker et al., 2011a), and deductive reasoning when able. MS/MS data was annotated by comparing resultant *m/z* peaks to the predicted values for fragment ions with up to 3 bond breaks from all possible parent structures using GlycoWorkbench (Damerell et al., 2012). Brain protein glycans were grouped into different categories based on shared components, such as monosaccharide composition, antennarity, etc., and the summed abundance of each category was compared across brain regions and sexes. All glycan structures are presented according to the Symbol Nomenclature for Glycans (SNFG) guidelines (Neelamegham et al., 2019; Varki et al., 2015) and were drawn using the GlycoGlyph online application (Mehta and Cummings, 2020).

#### Lectin Blotting

Brain lysate from the cortex and cerebellum of male mice, were precleared using magnetic streptavidin beads (New England Biolabs, #S1420S) at a 1:2 ratio of μg protein to μL washed beads to decrease background binding resulting from high levels of biotin-bound carboxylases in the brain. After 1-hour incubation at room temperature, beads and biotin-bound proteins were precipitated using a magnetic tube rack, and the supernatant was removed for further analysis. PNGase F sensitivity was determined by incubation of 100 μg protein with 5 μL PNGase F (New England Biolabs, #P0704S) at 37°C for 1 hour. Lysates were prepared with 4X Sample Loading Buffer (Li-COR, 928-40004) with 10% v/v β-mercaptoethanol, and denatured for 10 minutes at 95°C. For each gel, 15 μg protein was loaded per well (NuPAGE 4 to 12% Bis-Tris, 1.0 mm, Mini Protein Gel, 12-well, ThermoFisher, NP0322). In addition to 2 μL Chameleon Duo Pre-Stained Protein Ladder (LiCOR, 928-60000), 50 μg of human plasma was loaded as a positive control; plasma is ~60% is non-glycosylated albumin, thus ~20 μg plasma glycoprotein per lane. Gels were run using the MiniProtean Tetra Electrophoresis System (BioRAD, 1658004) at 140 mV for 1 hour. Proteins were transferred to nitrocellulose membranes (ThermoFisher, IB23003) using the iBlot Dry Blotting System (ThermoFisher, IB1001). Membranes were then incubated in 5% BSA in TBS-Tween 0.1% for 1 hour, followed by incubation with biotinylated lectins (Vector Labs: AAL B-1395, SNA B-1305, GNL B-1245, PHA-E B-1125, RCA B-1085, Con A B-1105) at a 1:1,000 dilution (1:20,000 for Con A) and 1:2,000 dilution of mouse anti-actin antibody (Abcam, ab8226) in 5% BSA in TBS-Tween 0.1%, overnight at 4°C on a rocking platform shaker. Membranes were washed three times in TBS-Tween 0.1% for 5 minutes, and then incubated with fluorescent conjugated streptavidin IRDye 800CW (LiCOR, 926-32230) and Goat anti-Mouse IgG IRDye 680RD (LiCOR, 925-68070) at 1:25,000 dilution in 5% BSA in TBS-Tween 0.1% for 30 minutes protected from light. Membranes were again washed three times in TBS-Tween 0.1% for 5 minutes and imaged using a LiCOR Odyssey CLx Imaging System and analyzed using LiCOR Image Studio Software. An identical unprobed membrane was incubated with Revert 700 Total Protein Stain (LiCOR, 926-11011) according to manufacturer’s protocol. A comprehensive characterization of biotinylated lectin binding specificity by glycan microarray can be found on the National Center for Functional Glycomics website (www.ncfg.hms.harvard.edu).

#### RNA Sequencing

Using the contralateral hemisphere of 4 male mouse brains used in glycomics and lectin blotting experiments, RNA from snap frozen cortex and cerebellum was purified using the RNeasy Lipid Tissue Mini Kit (QIAGEN, 74804) per manufacturer’s protocol. RNA-seq libraries were prepared from total RNA using polyA selection followed by the NEBNext Ultra II Directional RNA Library Prep Kit protocol (New England Biolabs, E7760S). Sequencing was performed on Illumina HiSeq 2500 instrument resulting in approximately 30 million of 50 bp reads per sample. Sequencing reads were mapped in a splice-aware fashion to the mouse reference transcriptome (mm9 assembly) using STAR (Dobin et al., 2013). Read counts over transcripts were calculated using HTSeq based on the Ensembl annotation for GRCm37/mm9 assembly and presented as Transcripts Per Million (TPM) (Anders et al., 2015). A subset of 269 known glycosyltransferases, glycosylhydrolases, sulfotransferases, and glycan-related genes was created, and differences in expression level between cortex and cerebellum was performed as described below.

#### Human RNA comparison and FUMA analysis

Performed utilizing publicly available gene expression data from the Genotype-Tissue Expression (GTEx) Portal, Version 8 (https://gtexportal.org). 354 known glycosyltransferases, glycosylhydrolases, sulfotransferases, and glycan-related genes IDs from humans were used as input into the GENE2FUNC platform of FUMA, which utilizes the GTEx v8 data of both 30 general tissue types, with all brain regions summarized as one tissue type, and 54 specific tissue types that includes 13 individual brain regions. Data is presented alphabetically, with significantly enriched differentially expressed gene sets presented in red after automatic *Bonferroni* correction with corrected p < 0.05.

### QUANTIFICATION AND STATISTICAL ANALYSIS

For glycomic analyses, statistical analysis of individual and groups of glycans was performed with Microsoft Excel Version 16.35. The abundance of individual glycans and glycan classes were compared between brain regions using single factor ANOVAs. Unpaired t-tests assuming unequal variance were performed for sex comparisons of individual N-glycans and glycan classes from cortex, cerebellum. Significance thresholds for ANOVAs and t-tests were applied at p < 0.05 (*), p < 0.01 (**), and p < 0.001 (***). A simple regression was performed between O-glycans modified with NeuAc or Fuc using GraphPad Prism v8.4.2. The EdgeR method was used for differential expression analysis of RNAseq data with gene cutoffs of 2-fold change in expression value and false discovery rates (FDR) below 0.05 as previously described (Robinson et al., 2010). Global glycosylation gene regulation in humans was analyzed using the FUMA GWAS GENE2FUC online tool, which identified significantly up- or down-regulated differentially expressed gene sets across human tissue types with a Bonferroni corrected p value < 0.05.

## Supplementary Files

**Supplementary data is presented in attached documents**

**Figs S1-S6 and Tables S1-S7**

**Table S8. RNA expression levels of glycosylation genes detected in cortex and**

**cerebellum**

**Table S9. Glycosylation-related genes included in FUMA analysis**

**File S1:** Original FUMA heat map of 354 glycosylation-related gene expression across tissues.

