## Supplementary Material for "The restricted nature of protein glycosylation in the mammalian brain"

**Mealer, et al,. 2020.**

**<u>Supplementary Figures</u>**

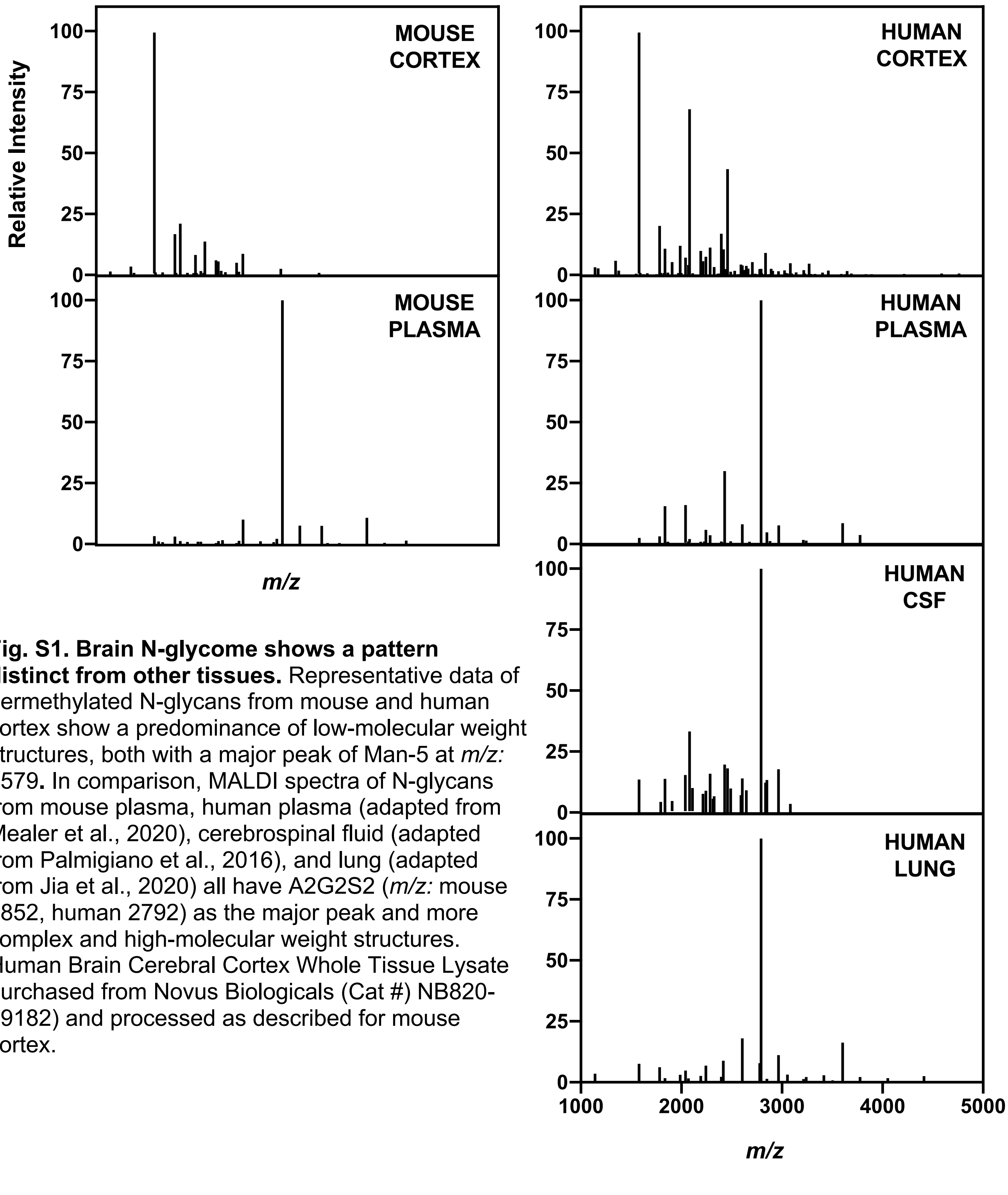


**Fig. S1. Brain N-glycome shows a pattern distinct from other tissues.** Representative data of permethylated N-glycans from mouse and human cortex show a predominance of low-molecular weight structures, both with a major peak of Man-5 at *m/z:* 1579**.** In comparison, MALDI spectra of N-glycans from mouse plasma, human plasma (adapted from Mealer et al., 2020), cerebrospinal fluid (adapted from Palmigiano et al., 2016), and lung (adapted from Jia et al., 2020) all have A2G2S2 (*m/z:* mouse 2852, human 2792) as the major peak and more complex and high-molecular weight structures. Human Brain Cerebral Cortex Whole Tissue Lysate purchased from Novus Biologicals (Cat #) NB820-59182) and processed as described for mouse cortex.

**The restricted nature of protein glycosylation in the mammalian brain**

**Mealer, et al,. 2020.**

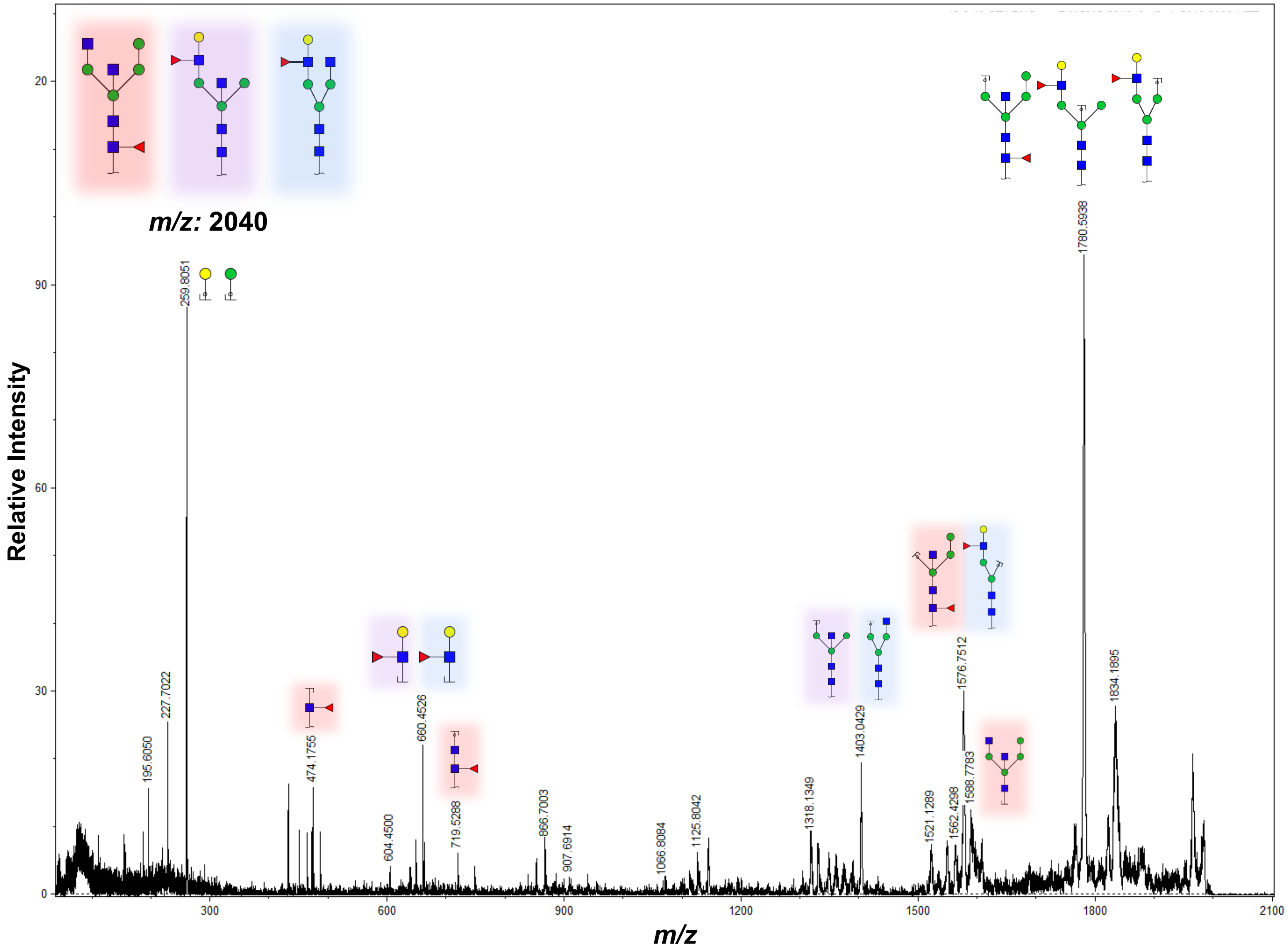


**Fig. S2. MS/MS analysis of *m/z:* 2040.** Fragmentation of the MALDI peak at *m/z:* 2040 revealed a mixture of glycan structures, including the hybrid structure $FA1BH_4$ (shown in red), due to the presence of fragment ions such as m/z: 1588, 719 and 474. Complex N-glycans (shown in purple and blue) present at this mass may account for the remaining signal after Endo H treatment. Related to Fig. 2.

**The restricted nature of protein glycosylation in the mammalian brain**

**Mealer, et al,. 2020.**

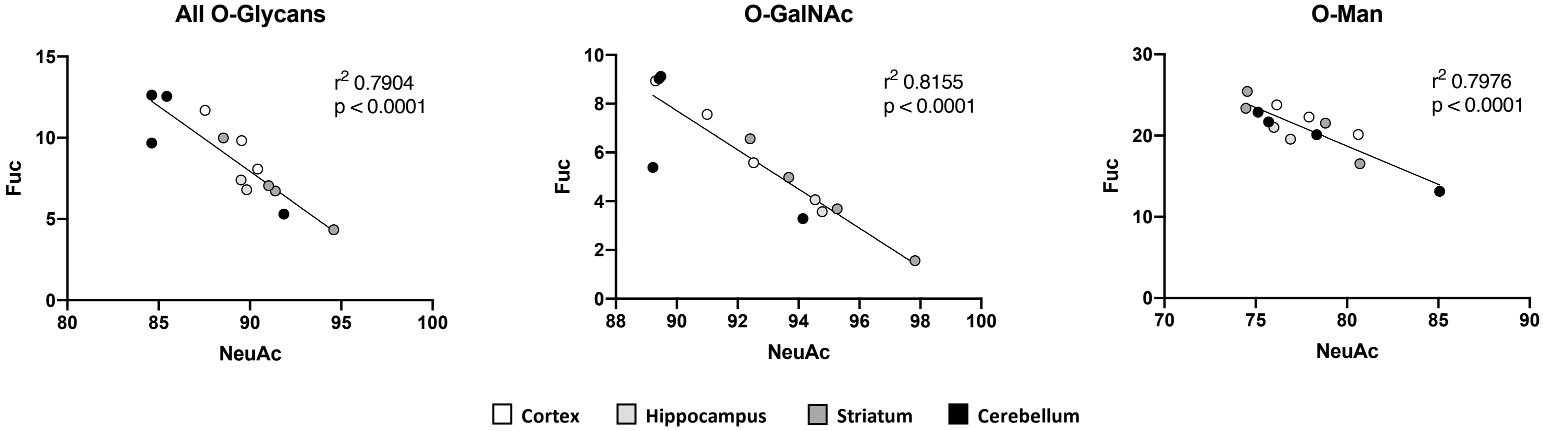


**Fig. S3. Negative correlation between fucose and NeuAc modifications on O-glycans in the brain.** Regression analysis demonstrated that the abundance of NeuAc-containing O-glycans is negatively correlated with the amount of fucose-containing O-glycans within a sample, suggesting competition between these modifications. This trend is observed in total O-glycans (A), O-GalNAc-type (B), and O-mannose-type glycans (C). Each data point represents a brain region from each mouse, color coded by region. Related to Fig. 3.

**The restricted nature of protein glycosylation in the mammalian brain**

**Mealer, et al,. 2020.**

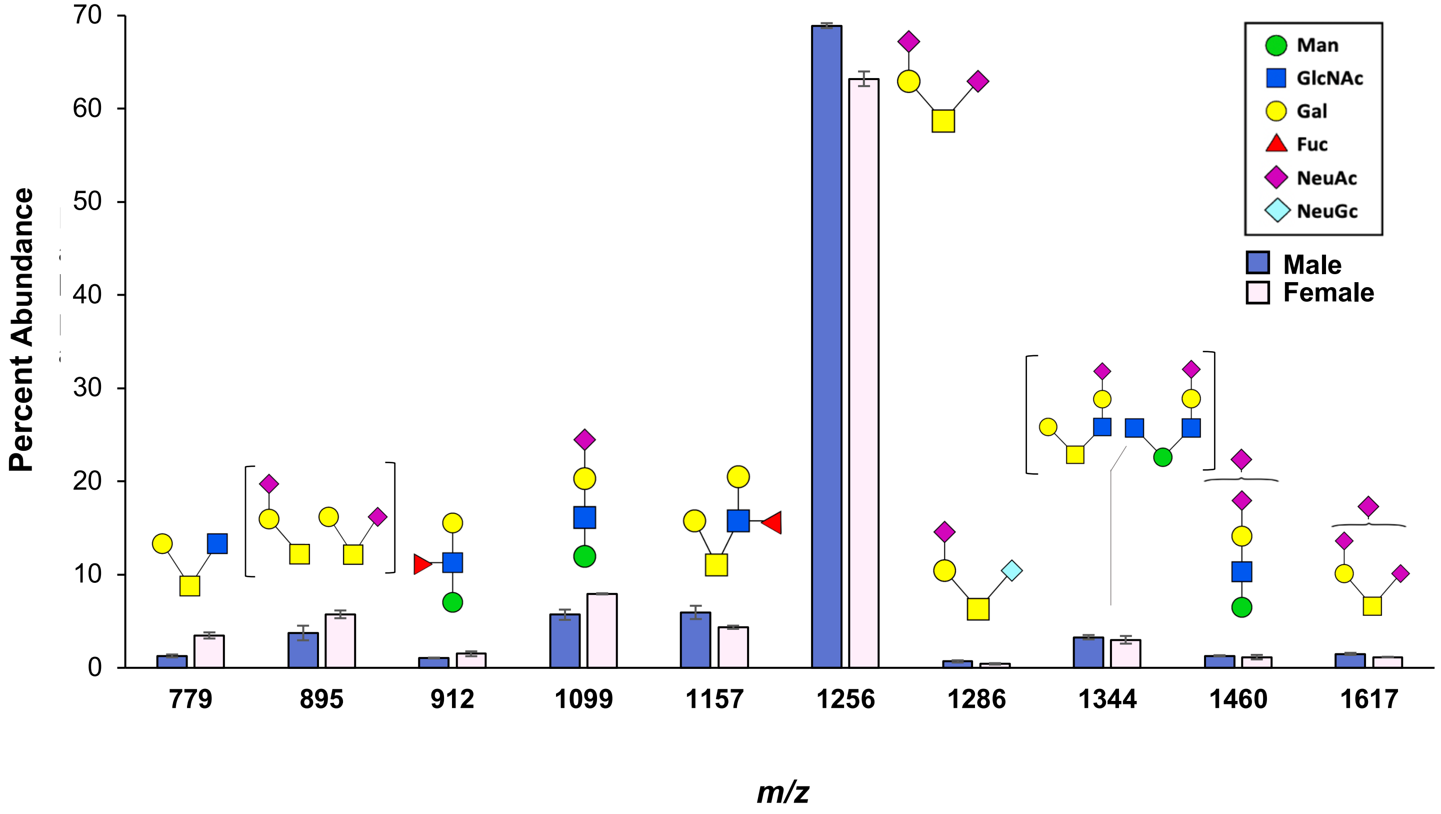


**Fig. S4. Sex-specific variation in protein O-glycosylation.** Comparison of the 10 most abundant protein O-glycans in the brain reveals minor variation between sexes, though limited sample size prevents statistical analysis (Male=3, Female=2). Data presented as mean percent abundance of each peak. Corresponding glycan structures are shown above each assigned peak. Related to Fig. 4.

**The restricted nature of protein glycosylation in the mammalian brain**

**Mealer, et al,. 2020.**

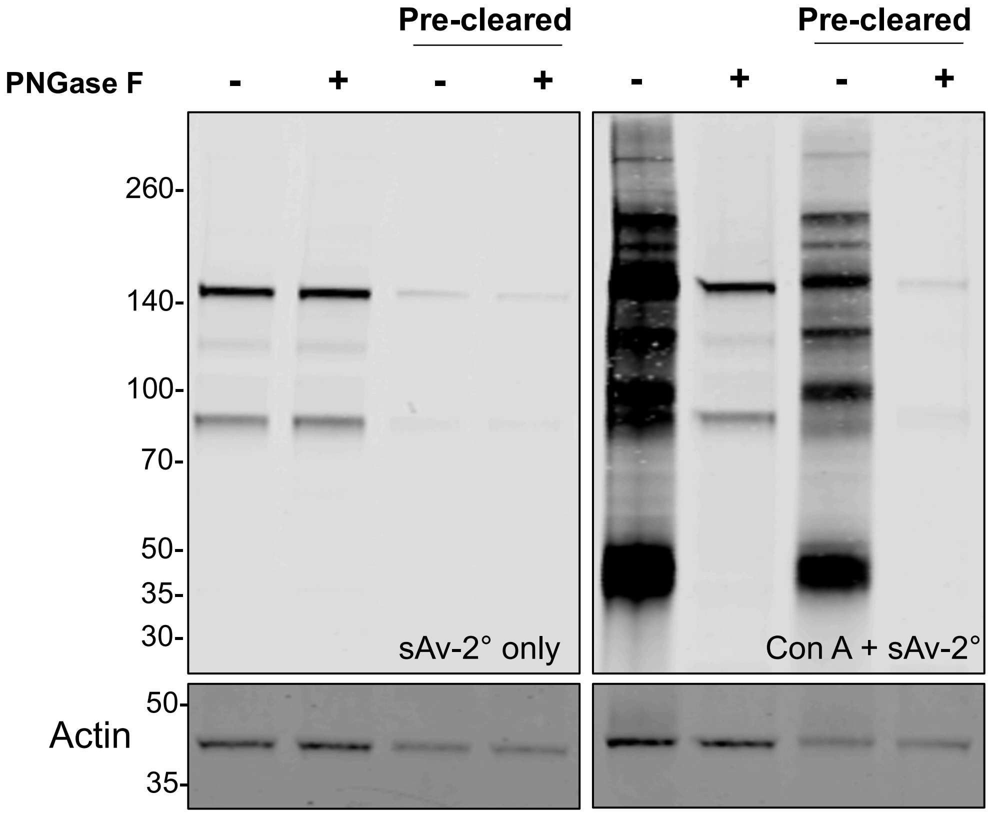


**Fig. S5: Streptavidin bead pre-clearing of brain lysate reduced non-specific binding to biotin-bound proteins.** Protein from mouse frontal cortex was incubated with or without magnetic streptavidin beads for 1 hour, followed by PNGase F digestion. Left panel was incubated with fluorescent streptavidin secondary alone (Licor sAv-800λ); right panel was incubated with primary biotinylated Con A followed by fluorescently labeled streptavidin secondary. Related to Fig. 5.

**The restricted nature of protein glycosylation in the mammalian brain**

**Mealer, et al,. 2020.**

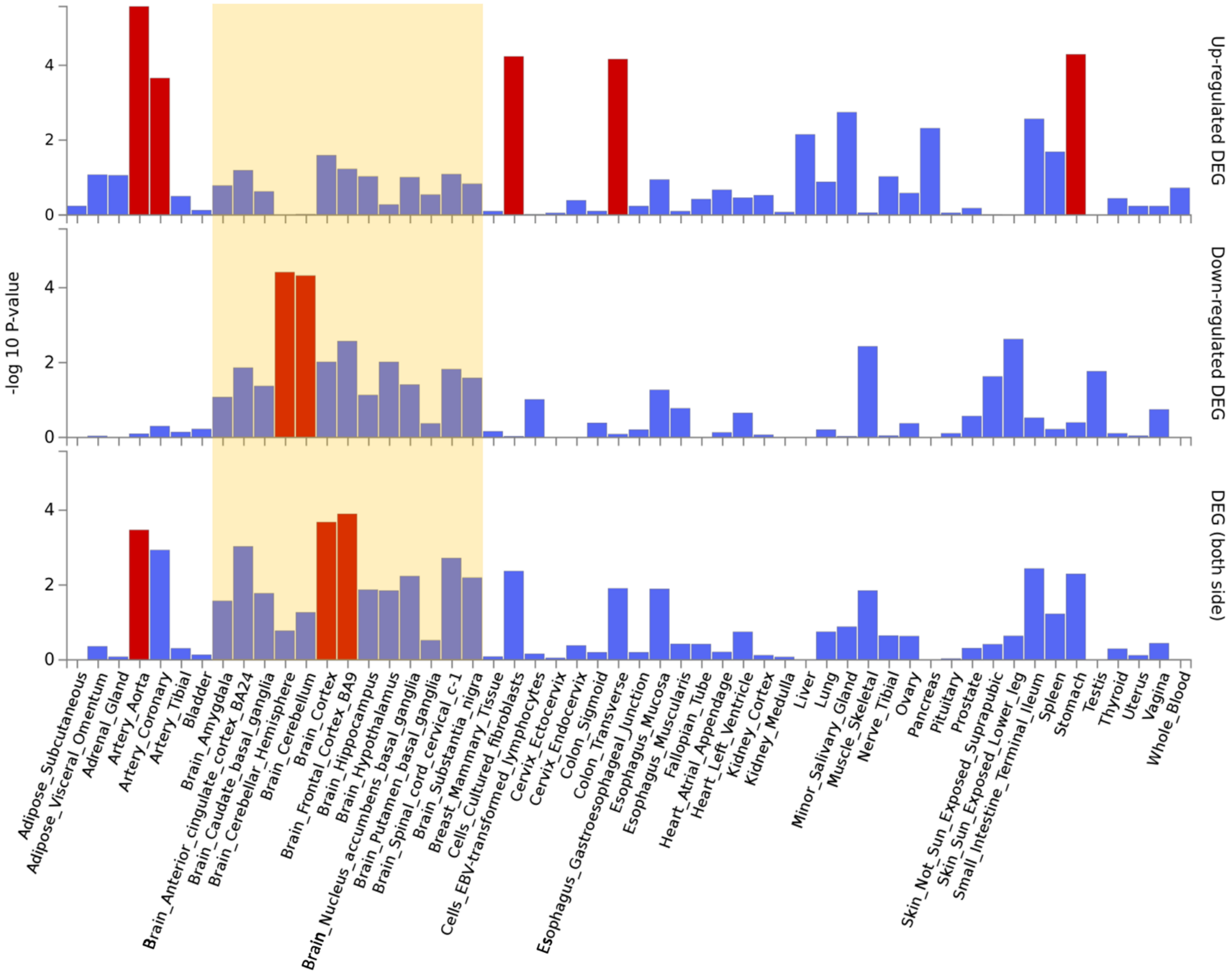


**Fig. S6: FUMA analysis revealed similar pattern of glycosylation gene regulation across brain regions.** Human tissue specific analysis shows some regional differences across 13 independent brain regions, but most show an overall pattern of downregulation, with significantly enriched DEG sets ($P_{bon}$ < 0.05) highlighted in red. Related to Fig. 7

### The restricted nature of protein glycosylation in the mammalian brain

### Mealer, et al,. 2020.

| **N-Glycans** | **Percent Abundance** | | | |
|---|---|---|---|---|
| ***m/z*** | CTX | HIP | STR | CBLM |
| **1141.62** | 0.654 | 0.554 | 0.643 | 0.286 |
| **1171.64** | 0.223 | 0.189 | 0.218 | 0.100 |
| **1345.75** | 1.569 | 1.180 | 1.101 | 0.452 |
| **1375.77** | 0.389 | 0.400 | 0.368 | 0.197 |
| **1416.80** | 0.107 | 0.119 | 0.113 | 0.068 |
| **1549.88** | 0.252 | 0.264 | 0.269 | 0.135 |
| **1579.89** | 45.857 | 46.105 | 45.096 | 37.127 |
| **1590.91** | 0.503 | 0.469 | 0.369 | 0.170 |
| **1620.93** | 0.142 | 0.153 | 0.146 | 0.118 |
| **1661.95** | 0.488 | 0.501 | 0.400 | 0.189 |
| **1754.01** | 0.174 | 0.206 | 0.178 | 0.137 |
| **1784.01** | 7.691 | 6.841 | 7.200 | 13.441 |
| **1795.03** | 0.406 | 0.442 | 0.426 | 0.235 |
| **1825.05** | 0.312 | 0.343 | 0.363 | 0.339 |
| **1836.05** | 9.627 | 9.802 | 9.132 | 3.739 |
| **1866.05** | 0.147 | 0.168 | 0.150 | 0.165 |
| **1907.11** | 0.416 | 0.464 | 0.390 | 0.632 |
| **1969.13** | 0.358 | 0.239 | 0.220 | 0.046 |
| **1988.13** | 3.761 | 3.045 | 3.413 | 4.555 |
| **1999.15** | 0.355 | 0.419 | 0.470 | 0.347 |
| **2040.17** | 0.745 | 0.907 | 0.803 | 0.889 |
| **2070.18** | 0.408 | 0.473 | 0.523 | 1.315 |
| **2081.20** | 6.296 | 8.394 | 7.929 | 12.988 |
| **2111.22** | 0.045 | 0.077 | 0.065 | 0.084 |
| **2192.24** | 2.738 | 2.490 | 2.539 | 2.667 |
| **2203.26** | 0.246 | 0.261 | 0.269 | 0.435 |
| **2214.26** | 2.547 | 1.946 | 2.084 | 0.611 |
| **2244.28** | 0.761 | 1.029 | 1.120 | 1.967 |
| **2285.31** | 0.517 | 0.648 | 0.537 | 0.693 |
| **2326.34** | 0.103 | 0.088 | 0.110 | 0.340 |
| **2360.35** | 0.063 | 0.099 | 0.090 | 0.079 |
| **2377.36** | 0.117 | 0.162 | 0.165 | 0.143 |
| **2390.36** | 0.091 | 0.113 | 0.128 | 0.178 |
| **2396.35** | 2.302 | 2.073 | 2.328 | 3.205 |
| **2418.37** | 0.614 | 0.490 | 0.686 | 0.603 |
| **2431.39** | 0.066 | 0.067 | 0.087 | 0.087 |
| **2448.40** | 0.215 | 0.196 | 0.281 | 0.345 |
| **2459.40** | 3.972 | 3.859 | 4.310 | 5.385 |
| **2489.42** | 0.061 | 0.083 | 0.072 | 0.084 |
| **2530.44** | 0.090 | 0.080 | 0.088 | 0.156 |
| **2547.42** | 0.028 | 0.048 | 0.029 | 0.069 |
| **2564.46** | 0.048 | 0.080 | 0.079 | 0.086 |
| **2592.47** | 0.296 | 0.228 | 0.232 | 0.127 |
| **2600.47** | 0.039 | 0.032 | 0.048 | 0.049 |
| **2605.48** | 0.086 | 0.105 | 0.115 | 0.121 |
| **2622.49** | 0.146 | 0.136 | 0.206 | 0.195 |
| **2635.50** | 0.078 | 0.083 | 0.101 | 0.128 |
| **2646.50** | 0.167 | 0.210 | 0.217 | 0.381 |
| **2663.51** | 0.164 | 0.156 | 0.161 | 0.184 |
| **2704.53** | 0.304 | 0.201 | 0.303 | 0.593 |
| **2762.57** | 0.028 | 0.039 | 0.035 | 0.022 |
| **2779.57** | 0.110 | 0.114 | 0.121 | 0.086 |
| **2796.59** | 0.053 | 0.056 | 0.073 | 0.058 |
| **2809.59** | 0.038 | 0.047 | 0.057 | 0.057 |
| **2837.60** | 1.172 | 0.854 | 0.997 | 0.752 |

| **O-Glycans** | **Percent Abundance** | | | |
|---|---|---|---|---|
| ***m/z*** | CTX | HIP | STR | CBLM |
| **534.30** | 0.582 | 1.377 | 0.802 | 0.540 |
| **691.41** | 0.247 | 0.255 | 0.181 | 0.530 |
| **779.46** | 1.280 | 0.988 | 0.789 | 2.439 |
| **895.50** | 3.739 | 6.551 | 3.662 | 7.197 |
| **912.51** | 1.055 | 2.579 | 1.395 | 4.639 |
| **925.53** | 0.128 | 0.177 | 0.150 | 0.360 |
| **983.59** | 0.389 | 0.278 | 0.290 | 0.391 |
| **1069.63** | 0.237 | 0.084 | 0.254 | 0.794 |
| **1099.62** | 5.698 | 11.386 | 7.210 | 18.380 |
| **1129.65** | 0.218 | 0.814 | 0.360 | 0.473 |
| **1157.66** | 5.947 | 2.885 | 3.176 | 4.032 |
| **1256.69** | 68.923 | 64.266 | 68.654 | 55.196 |
| **1286.70** | 0.690 | 0.432 | 1.130 | 1.068 |
| **1316.72** | 0.176 | 0.165 | 0.226 | 0.145 |
| **1344.75** | 3.285 | 1.694 | 2.406 | 0.744 |
| **1361.76** | 0.445 | 0.412 | 0.672 | 0.128 |
| **1460.80** | 1.293 | 1.403 | 1.399 | 0.732 |
| **1518.84** | 1.330 | 0.305 | 0.595 | 0.261 |
| **1535.86** | 0.526 | 0.513 | 0.519 | 0.104 |
| **1548.85** | 0.551 | 0.612 | 0.790 | 0.159 |
| **1589.87** | 0.077 | 0.134 | 0.232 | 0.109 |
| **1617.88** | 1.498 | 1.281 | 3.053 | 0.940 |
| **1705.95** | 0.772 | 0.277 | 0.601 | 0.322 |
| **1722.97** | 0.328 | 0.331 | 0.420 | 0.087 |
| **1910.08** | 0.491 | 0.731 | 0.853 | 0.158 |
| **1979.12** | 0.096 | 0.069 | 0.181 | 0.072 |

| | | | | |
|---|---|---|---|---|
| **2867.59** | 0.024 | 0.024 | 0.030 | 0.037 |
| **2891.64** | 0.076 | 0.088 | 0.106 | 0.221 |
| **2908.64** | 0.057 | 0.053 | 0.067 | 0.108 |
| **2925.62** | 0.031 | 0.040 | 0.034 | 0.037 |
| **2966.68** | 0.053 | 0.075 | 0.073 | 0.080 |
| **2983.67** | 0.013 | 0.018 | 0.027 | 0.030 |
| **2996.70** | 0.020 | 0.021 | 0.025 | 0.032 |
| **3024.70** | 0.134 | 0.130 | 0.136 | 0.133 |
| **3041.72** | 0.049 | 0.058 | 0.060 | 0.055 |
| **3054.71** | 0.016 | 0.022 | 0.025 | 0.024 |
| **3082.74** | 0.153 | 0.129 | 0.191 | 0.271 |
| **3095.76** | 0.014 | 0.022 | 0.022 | 0.039 |
| **3140.77** | 0.039 | 0.042 | 0.050 | 0.032 |
| **3215.81** | 0.383 | 0.536 | 0.391 | 0.124 |
| **3228.83** | 0.025 | 0.032 | 0.033 | 0.030 |
| **3269.84** | 0.104 | 0.110 | 0.146 | 0.190 |
| **3286.84** | 0.029 | 0.032 | 0.042 | 0.063 |
| **3327.88** | 0.031 | 0.036 | 0.044 | 0.049 |
| **3402.91** | 0.147 | 0.227 | 0.184 | 0.081 |
| **3456.96** | 0.013 | 0.020 | 0.023 | 0.037 |
| **3460.95** | 0.064 | 0.058 | 0.083 | 0.098 |
| **3473.98** | 0.011 | 0.016 | 0.019 | 0.028 |
| **3515.01** | 0.011 | 0.007 | 0.014 | 0.018 |
| **3590.03** | 0.034 | 0.051 | 0.051 | 0.037 |
| **3631.06** | 0.014 | 0.020 | 0.020 | 0.019 |
| **3648.06** | 0.069 | 0.059 | 0.083 | 0.076 |
| **3764.13** | 0.014 | 0.018 | 0.026 | 0.023 |
| **3777.15** | 0.009 | 0.019 | 0.019 | 0.021 |
| **3835.18** | 0.020 | 0.020 | 0.026 | 0.029 |
| **3852.16** | 0.008 | 0.014 | 0.019 | 0.026 |
| **3893.22** | 0.013 | 0.009 | 0.020 | 0.025 |
| **4009.29** | 0.014 | 0.014 | 0.021 | 0.020 |
| **4026.30** | 0.059 | 0.089 | 0.115 | 0.113 |
| **4039.30** | 0.007 | 0.009 | 0.015 | 0.018 |
| **4080.35** | 0.005 | 0.003 | 0.009 | 0.015 |
| **4213.44** | 0.028 | 0.033 | 0.059 | 0.063 |
| **4400.57** | 0.009 | 0.009 | 0.018 | 0.029 |
| **4574.70** | 0.006 | 0.004 | 0.008 | 0.009 |
| **4587.69** | 0.002 | 0.002 | 0.005 | 0.009 |
| **4649.74** | 0.003 | 0.005 | 0.006 | 0.008 |

**Table S1. Individual protein glycan abundance across brain regions.** Data presented as mean percent abundance. For N-glycans, n=6 for each brain region. For O-glycans, CTX=3, HIP=2, STR=4, CBLM=4. Related to Fig. 1 and Fig. 3.

#### The restricted nature of protein glycosylation in the mammalian brain

#### Mealer, et al,. 2020.

| Glycan Name | *m/z* | GlcNAc | Mannose | Galactose | Fucose | NeuAc | Pauci | High-Man | Hybrid | Bisected | Antenna |
|---|---|---|---|---|---|---|---|---|---|---|---|
| F-Man-2 | 1141.6 | 2 | 2 | 0 | 1 | 0 | 1 | 0 | 0 | 0 | 0 |
| Man-3 | 1171.1 | 2 | 3 | 0 | 0 | 0 | 1 | 0 | 0 | 0 | 0 |
| F-Man-3 | 1345.8 | 2 | 3 | 0 | 1 | 0 | 1 | 0 | 0 | 0 | 0 |
| Man-4 | 1375.8 | 2 | 4 | 0 | 0 | 0 | 1 | 0 | 0 | 0 | 0 |
| A1 | 1416.8 | 3 | 3 | 0 | 0 | 0 | 0 | 0 | 0 | 0 | 1 |
| F-Man-4 | 1549.9 | 2 | 4 | 0 | 1 | 0 | 1 | 0 | 0 | 0 | 0 |
| Man-5 | 1579.9 | 2 | 5 | 0 | 0 | 0 | 0 | 1 | 0 | 0 | 0 |
| FA1 | 1590.9 | 3 | 3 | 0 | 1 | 0 | 0 | 0 | 0 | 0 | 1 |
| A1H4 | 1620.9 | 3 | 4 | 0 | 0 | 0 | 0 | 0 | 1 | 0 | 1 |
| A1B | 1661.9 | 4 | 3 | 0 | 0 | 0 | 0 | 0 | 0 | 1 | 1 |
| F-Man-5 | 1754.0 | 2 | 5 | 0 | 1 | 0 | 0 | 1 | 0 | 0 | 0 |
| Man-6 | 1784.0 | 2 | 6 | 0 | 0 | 0 | 0 | 1 | 0 | 0 | 0 |
| FA1H4 | 1795.0 | 3 | 4 | 0 | 1 | 0 | 0 | 0 | 1 | 0 | 1 |
| A1H5 | 1825.0 | 3 | 5 | 0 | 0 | 0 | 0 | 0 | 1 | 0 | 1 |
| FA1B | 1836.0 | 4 | 3 | 0 | 1 | 0 | 0 | 0 | 0 | 1 | 1 |
| A1BH4 | 1866.1 | 4 | 4 | 0 | 0 | 0 | 0 | 0 | 1 | 1 | 1 |
| A2B | 1907.1 | 5 | 3 | 0 | 0 | 0 | 0 | 0 | 0 | 1 | 2 |
| F2A1G1 | 1969.1 | 3 | 3 | 1 | 2 | 0 | 0 | 0 | 0 | 0 | 1 |
| Man-7 | 1988.1 | 2 | 7 | 0 | 0 | 0 | 0 | 1 | 0 | 0 | 0 |
| FA1H | 1999.2 | 3 | 5 | 0 | 1 | 0 | 0 | 0 | 1 | 0 | 1 |
| FA1BH4 | 2040.2 | 4 | 4 | 0 | 1 | 0 | 0 | 0 | 1 | 1 | 1 |
| A1BH5 | 2070.2 | 4 | 5 | 0 | 0 | 0 | 0 | 0 | 1 | 1 | 1 |
| FA2B | 2081.2 | 5 | 3 | 0 | 1 | 0 | 0 | 0 | 0 | 1 | 2 |
| A2BH4 | 2111.2 | 5 | 4 | 0 | 0 | 0 | 0 | 0 | 1 | 1 | 2 |
| Man-8 | 2192.2 | 2 | 8 | 0 | 0 | 0 | 0 | 1 | 0 | 0 | 0 |
| FA1G1H5 | 2203.2 | 3 | 5 | 1 | 1 | 0 | 0 | 0 | 1 | 0 | 1 |
| F2A1G1B | 2214.3 | 4 | 3 | 1 | 2 | 0 | 0 | 0 | 0 | 1 | 1 |
| FA1BH5 | 2244.3 | 4 | 5 | 0 | 1 | 0 | 0 | 0 | 1 | 1 | 1 |
| FA2BG1 | 2285.3 | 5 | 3 | 1 | 1 | 0 | 0 | 0 | 0 | 1 | 2 |
| FA3B | 2326.3 | 6 | 3 | 0 | 1 | 0 | 0 | 0 | 0 | 1 | 3 |
| FA1G1S1H4 | 2360.3 | 3 | 4 | 1 | 1 | 1 | 0 | 0 | 1 | 0 | 1 |
| F2A1G1H5 | 2377.3 | 3 | 5 | 1 | 2 | 0 | 0 | 0 | 1 | 0 | 1 |
| A1G1S1H5 | 2390.3 | 3 | 5 | 1 | 0 | 1 | 0 | 0 | 1 | 0 | 1 |
| Man-9 | 2396.3 | 2 | 9 | 0 | 0 | 0 | 0 | 1 | 0 | 0 | 0 |
| F2A1G1BH4 | 2418.4 | 4 | 4 | 1 | 2 | 0 | 0 | 0 | 1 | 1 | 1 |
| A1G1S1BH4 | 2431.4 | 4 | 4 | 1 | 0 | 1 | 0 | 0 | 1 | 1 | 1 |
| FAG1BH5 | 2448.3 | 4 | 5 | 1 | 1 | 0 | 0 | 0 | 1 | 1 | 1 |
| F2A1G1B | 2459.4 | 5 | 3 | 1 | 2 | 0 | 0 | 0 | 0 | 1 | 2 |
| FA2G1BH4 | 2489.4 | 5 | 4 | 1 | 1 | 0 | 0 | 0 | 1 | 1 | 2 |
| FA3G1B | 2530.4 | 6 | 3 | 1 | 1 | 0 | 0 | 0 | 0 | 1 | 3 |
| A1G1S2H4 | 2547.4 | 3 | 4 | 1 | 0 | 2 | 0 | 0 | 1 | 0 | 1 |
| FA1G1S1H5 | 2564.4 | 3 | 5 | 1 | 1 | 1 | 0 | 0 | 1 | 0 | 1 |
| F3A2G2 | 2592.5 | 4 | 3 | 2 | 3 | 0 | 0 | 0 | 0 | 0 | 2 |
| Man-9-G | 2600.5 | 2 | 9 | 0 | 0 | 0 | 0 | 1 | 0 | 0 | 0 |
| FA1G1S1BH4 | 2605.5 | 4 | 4 | 1 | 1 | 1 | 0 | 0 | 1 | 1 | 1 |
| F2A1G1BH5 | 2622.5 | 4 | 5 | 1 | 2 | 0 | 0 | 0 | 1 | 1 | 1 |
| A1G1S1BH5 | 2635.5 | 4 | 5 | 1 | 0 | 1 | 0 | 0 | 1 | 1 | 1 |
| FA2G1S1B | 2646.5 | 5 | 3 | 1 | 1 | 1 | 0 | 0 | 0 | 1 | 2 |
| F2A2G2B | 2663.5 | 5 | 3 | 2 | 2 | 0 | 0 | 0 | 0 | 1 | 2 |
| F2A3G1B | 2704.5 | 6 | 3 | 1 | 2 | 0 | 0 | 0 | 0 | 1 | 3 |
| FA1G1S2B | 2762.5 | 4 | 3 | 1 | 1 | 2 | 0 | 0 | 0 | 1 | 1 |
| F2A1G1S1BH4 | 2779.6 | 4 | 4 | 1 | 2 | 1 | 0 | 0 | 1 | 1 | 1 |
| F3A1G1BH5 | 2796.6 | 4 | 5 | 1 | 3 | 0 | 0 | 0 | 1 | 1 | 1 |
| FA1G1S1BH5 | 2809.6 | 4 | 5 | 1 | 1 | 1 | 0 | 0 | 1 | 1 | 1 |

| | | | | | | | | | | | |
|---|---|---|---|---|---|---|---|---|---|---|---|
| F3A2G2B | 2837.6 | 5 | 3 | 2 | 3 | 0 | 0 | 0 | 0 | 1 | 2 |
| F2A2G2BH4 | 2867.6 | 5 | 4 | 2 | 2 | 0 | 0 | 0 | 0 | 1 | 2 |
| FA3G1S1B | 2891.6 | 6 | 3 | 1 | 1 | 1 | 0 | 0 | 0 | 1 | 3 |
| F2A3G2B | 2908.6 | 6 | 3 | 2 | 2 | 0 | 0 | 0 | 0 | 1 | 3 |
| FA1G1S2H5 | 2925.6 | 3 | 5 | 1 | 1 | 2 | 0 | 0 | 1 | 0 | 1 |
| FA2G2S2 | 2966.6 | 4 | 3 | 2 | 1 | 2 | 0 | 0 | 0 | 0 | 2 |
| F2A1G1S1BH5 | 2983.6 | 4 | 5 | 1 | 2 | 1 | 0 | 0 | 1 | 1 | 1 |
| A1G1S2BH | 2996.6 | 4 | 5 | 1 | 0 | 2 | 0 | 0 | 1 | 1 | 1 |
| F2A2G2S1B | 3024.7 | 5 | 3 | 2 | 2 | 1 | 0 | 0 | 0 | 1 | 2 |
| F3A3G3 | 3041.7 | 5 | 3 | 3 | 3 | 0 | 0 | 0 | 0 | 0 | 3 |
| FA2G2S1BH4 | 3054.7 | 5 | 4 | 2 | 1 | 1 | 0 | 0 | 1 | 1 | 2 |
| F3A3G2B | 3082.7 | 6 | 3 | 2 | 3 | 0 | 0 | 0 | 0 | 1 | 3 |
| FA3G2S1B | 3095.7 | 6 | 3 | 2 | 1 | 1 | 0 | 0 | 0 | 1 | 3 |
| F2A2G2S2 | 3140.8 | 4 | 3 | 2 | 2 | 2 | 0 | 0 | 0 | 0 | 2 |
| F4A3G3 | 3215.8 | 5 | 3 | 3 | 4 | 0 | 0 | 0 | 0 | 0 | 3 |
| F2A3G3S1 | 3228.8 | 5 | 3 | 3 | 2 | 1 | 0 | 0 | 0 | 0 | 3 |
| F2A3G2S1B | 3269.8 | 6 | 3 | 2 | 2 | 1 | 0 | 0 | 0 | 1 | 3 |
| F3A3G3B | 3286.9 | 6 | 3 | 3 | 3 | 0 | 0 | 0 | 0 | 1 | 3 |
| FA2G2S3 | 3327.9 | 4 | 3 | 2 | 1 | 3 | 0 | 0 | 0 | 0 | 2 |
| F3A2G2S1 | 3402.9 | 4 | 3 | 2 | 3 | 1 | 0 | 0 | 0 | 0 | 2 |
| FA3G2S2B | 3456.9 | 6 | 3 | 2 | 1 | 2 | 0 | 0 | 0 | 1 | 3 |
| F4A3G3B | 3460.9 | 6 | 3 | 3 | 4 | 0 | 0 | 0 | 0 | 1 | 3 |
| F2A3G3S1B | 3473.9 | 6 | 3 | 3 | 2 | 1 | 0 | 0 | 0 | 1 | 3 |
| F2A4G2S1B | 3514.8 | 7 | 3 | 2 | 2 | 1 | 0 | 0 | 0 | 1 | 4 |
| F2A3G3S2 | 3590.0 | 5 | 3 | 3 | 2 | 2 | 0 | 0 | 0 | 0 | 3 |
| F2A3G2S2B | 3631.0 | 6 | 3 | 2 | 2 | 2 | 0 | 0 | 0 | 1 | 3 |
| F3A3G3S1B | 3648.0 | 6 | 3 | 3 | 3 | 1 | 0 | 0 | 0 | 1 | 3 |
| F3A3G3S2 | 3764.1 | 5 | 3 | 3 | 3 | 2 | 0 | 0 | 0 | 0 | 3 |
| FA3G3S3 | 3777.1 | 5 | 3 | 3 | 1 | 3 | 0 | 0 | 0 | 0 | 3 |
| F2A3G3S2B | 3835.2 | 6 | 3 | 3 | 2 | 2 | 0 | 0 | 0 | 1 | 3 |
| F3A4G4S1 | 3852.2 | 6 | 3 | 4 | 3 | 1 | 0 | 0 | 0 | 0 | 4 |
| F3A4G3S1B | 3893.2 | 7 | 3 | 3 | 3 | 1 | 0 | 0 | 0 | 1 | 4 |
| F3A3G3S2B | 4009.2 | 6 | 3 | 3 | 3 | 2 | 0 | 0 | 0 | 1 | 3 |
| F4A4G4S1 | 4026.3 | 6 | 3 | 4 | 4 | 1 | 0 | 0 | 0 | 0 | 4 |
| F2A4G4S2 | 4039.3 | 6 | 3 | 4 | 2 | 2 | 0 | 0 | 0 | 0 | 4 |
| F2A4G3S2B | 4080.4 | 7 | 3 | 3 | 2 | 2 | 0 | 0 | 0 | 1 | 4 |
| F3A4G4S2 | 4213.4 | 6 | 3 | 4 | 3 | 2 | 0 | 0 | 0 | 0 | 4 |
| F2A4G4S3 | 4400.5 | 6 | 3 | 4 | 2 | 3 | 0 | 0 | 0 | 0 | 4 |
| F3A4G4S3 | 4574.6 | 6 | 3 | 4 | 3 | 3 | 0 | 0 | 0 | 0 | 4 |
| FA4G4S4 | 4587.6 | 6 | 3 | 4 | 1 | 4 | 0 | 0 | 0 | 0 | 4 |
| F5A5G5S1 | 4649.6 | 7 | 3 | 5 | 5 | 1 | 0 | 0 | 0 | 0 | 5 |

**Table S2. Brain protein N-glycan structure, name, mass, and characteristics.** Related to Fig. 1 and Table 1.

**The restricted nature of protein glycosylation in the mammalian brain**

**Mealer, et al,. 2020.**

| Glycan Name | *m/z* | HexNAc | Hexose | NeuAc | NeuGc | All Sia | Fucose | Ser/Thr O-Man | Ser/Thr O-GalNAc | Ambiguous |
|---|---|---|---|---|---|---|---|---|---|---|
| HexNAcHex | 534.3 | 1 | 1 | 0 | 0 | 0 | 0 | 0 | 0 | 1 |
| HexNAcNeuAc | 691.4 | 1 | 0 | 1 | 0 | 1 | 0 | 0 | 1 | 0 |
| HexNAc2Hex | 779.4 | 2 | 1 | 0 | 0 | 0 | 0 | 0 | 1 | 0 |
| HexNAcHexNeuAc | 895.5 | 1 | 1 | 1 | 0 | 1 | 0 | 0 | 1 | 0 |
| HexNAcHex2Fuc | 912.5 | 1 | 2 | 0 | 0 | 0 | 1 | 1 | 0 | 0 |
| HexNAcHexNeuGc | 925.5 | 1 | 1 | 0 | 1 | 1 | 0 | 0 | 1 | 0 |
| HexNAc2Hex2 | 983.5 | 2 | 2 | 0 | 0 | 0 | 0 | 0 | 1 | 0 |
| HexNAcHexNeuAcFuc | 1069.6 | 1 | 1 | 1 | 0 | 1 | 1 | 0 | 1 | 0 |
| HexNAcHex2NeuAc | 1099.6 | 1 | 2 | 1 | 0 | 1 | 0 | 1 | 0 | 0 |
| HexNAcHexNeuGc | 1129.6 | 1 | 2 | 0 | 1 | 1 | 0 | 1 | 0 | 0 |
| HexNAc2Hex2Fuc | 1157.6 | 2 | 2 | 0 | 0 | 0 | 1 | 0 | 1 | 0 |
| HexNAcHexNeuAc2 | 1256.6 | 1 | 1 | 2 | 0 | 2 | 0 | 0 | 1 | 0 |
| HexNAcHexNeuAcNeuGc | 1286.7 | 1 | 1 | 1 | 1 | 2 | 0 | 0 | 1 | 0 |
| HexNAcHexNeuGc2 | 1316.7 | 1 | 1 | 0 | 2 | 2 | 0 | 0 | 1 | 0 |
| HexNAc2Hex2NeuAc | 1344.7 | 2 | 2 | 1 | 0 | 1 | 0 | 0 | 0 | 1 |
| HexNAc2Hex3Fuc | 1361.7 | 2 | 3 | 0 | 0 | 0 | 1 | 1 | 0 | 0 |
| HexNAcHex2NeuAc2 | 1460.8 | 1 | 2 | 2 | 0 | 2 | 0 | 1 | 0 | 0 |
| HexNAc2Hex2NeuAcFuc | 1518.8 | 2 | 2 | 1 | 0 | 1 | 1 | 0 | 0 | 1 |
| HexNAc2Hex3Fuc2 | 1535.8 | 2 | 3 | 0 | 0 | 0 | 2 | 1 | 0 | 0 |
| HexNAc2Hex3NeuAc | 1548.8 | 2 | 3 | 1 | 0 | 1 | 0 | 1 | 0 | 0 |
| HexNAc2Hex3NeuGc | 1589.8 | 2 | 3 | 0 | 1 | 1 | 0 | 1 | 0 | 0 |
| HexNAcHexNeuAc3 | 1617.8 | 1 | 1 | 3 | 0 | 3 | 0 | 0 | 1 | 0 |
| HexNAc2Hex2NeuAc2 | 1705.9 | 2 | 2 | 2 | 0 | 2 | 0 | 0 | 1 | 0 |
| HexNAc2Hex3NeuAcFuc | 1722.9 | 2 | 3 | 1 | 0 | 1 | 1 | 1 | 0 | 0 |
| HexNAc2Hex3NeuAc2 | 1910.0 | 2 | 3 | 2 | 0 | 2 | 0 | 1 | 0 | 0 |
| HexNAcHexNeuAc4 | 1979.1 | 1 | 1 | 4 | 0 | 4 | 0 | 0 | 1 | 0 |

**Table S3. Brain protein O-glycan structure, name, mass, and characteristics.** Related to Fig. 3 and Table 1.

### The restricted nature of protein glycosylation in the mammalian brain

### Mealer, et al,. 2020.

| | Plasma N-glycans | | | | Cortex N-glycans | | | Cerebellum N-glycans | | |
|---|---|---|---|---|---|---|---|---|---|---|
| *m/z* | Male | Female | p-value | *m/z* | Male | Female | p-value | Male | Female | p-value |
| 1579.9 | 0.35 | 0.31 | 0.76 | 1141.7 | 0.66 | 0.89 | 0.18 | 0.29 | 0.91 | 0.15 |
| 1621.1 | 0.20 | 0.19 | 0.88 | 1171.7 | 0.22 | 0.24 | 0.73 | 0.10 | 0.25 | 0.15 |
| 1662.1 | 0.14 | 0.17 | 0.63 | 1345.8 | 1.58 | 1.48 | 0.55 | 0.45 | 0.98 | 0.15 |
| 1784.0 | 0.50 | 0.42 | 0.77 | 1375.8 | 0.39 | 0.41 | 0.72 | 0.20 | 0.48 | **0.046*** |
| 1836.1 | 0.27 | 0.42 | 0.41 | 1416.8 | 0.11 | 0.19 | **0.006**** | 0.07 | 0.16 | **0.0002***** |
| 1907.3 | 0.28 | 0.16 | 0.17 | 1549.9 | 0.25 | 0.31 | **0.02*** | 0.14 | 0.25 | 0.11 |
| 1988.1 | 0.10 | 0.10 | 0.98 | 1579.9 | 46.09 | 45.56 | 0.83 | 37.17 | 34.47 | 0.41 |
| 2012.2 | 0.13 | 0.09 | 0.28 | 1590.9 | 0.51 | 0.60 | **0.008**** | 0.17 | 0.29 | **0.013*** |
| 2040.2 | 0.26 | 0.62 | 0.33 | 1620.9 | 0.14 | 0.21 | **0.003**** | 0.12 | 0.28 | **0.003**** |
| 2192.2 | 0.07 | 0.08 | 0.88 | 1662.0 | 0.49 | 0.59 | 0.31 | 0.19 | 0.23 | 0.07 |
| 2216.3 | 0.37 | 0.28 | 0.44 | 1754.0 | 0.18 | 0.21 | 0.29 | 0.14 | 0.24 | **0.008**** |
| 2257.3 | 0.54 | 0.36 | 0.07 | 1784.0 | 7.73 | 6.78 | 0.07 | 13.46 | 13.92 | 0.54 |
| 2396.3 | 0.11 | 0.14 | 0.59 | 1795.0 | 0.41 | 0.52 | 0.06 | 0.24 | 0.42 | **0.002**** |
| 2420.4 | 0.46 | 0.33 | 0.47 | 1825.1 | 0.31 | 0.40 | **0.02*** | 0.34 | 0.62 | **0.008**** |
| 2431.4 | 0.13 | 0.20 | 0.11 | 1836.1 | 9.68 | 9.81 | 0.84 | 3.74 | 3.22 | 0.09 |
| 2461.4 | 5.15 | 2.04 | 0.07 | 1866.1 | 0.15 | 0.17 | 0.11 | 0.16 | 0.30 | **0.03*** |
| 2635.5 | 0.49 | 1.15 | 0.09 | 1907.1 | 0.42 | 0.56 | **0.02*** | 0.63 | 0.74 | 0.11 |
| 2852.6 | 53.84 | 33.85 | **0.0008***** | 1969.1 | 0.36 | 0.41 | **0.04*** | 0.05 | 0.08 | **0.004**** |
| 2910.6 | 0.24 | 0.14 | 0.012* | 1988.1 | 3.78 | 3.16 | **0.03*** | 4.56 | 4.77 | 0.34 |
| 3026.7 | 6.85 | 38.34 | **0.00014***** | 1999.2 | 0.36 | 0.43 | 0.12 | 0.35 | 0.50 | **0.03*** |
| 3243.8 | 7.28 | 2.02 | **0.0003***** | 2040.2 | 0.75 | 0.91 | 0.12 | 0.89 | 1.15 | 0.10 |
| 3301.8 | 0.53 | 0.36 | **0.0019**** | 2070.2 | 0.41 | 0.43 | 0.76 | 1.32 | 1.95 | 0.07 |
| 3417.9 | 0.43 | 1.61 | **0.005**** | 2081.2 | 6.33 | 6.62 | 0.69 | 13.00 | 9.47 | **0.005**** |
| 3693.1 | 16.88 | 10.20 | **0.0013**** | 2111.2 | - | - | - | 0.08 | 0.10 | 0.33 |
| 3867.2 | 0.90 | 5.11 | **0.002**** | 2192.2 | 2.75 | 2.24 | **0.03*** | 2.67 | 3.14 | 0.13 |
| 4084.3 | 3.26 | 1.00 | **0.002**** | 2203.2 | 0.25 | 0.27 | 0.41 | 0.44 | 0.66 | **0.01*** |
| 4258.4 | 0.07 | 0.22 | **0.04*** | 2214.3 | 2.56 | 2.48 | 0.51 | 0.61 | 0.65 | 0.62 |
| 4475.5 | 0.08 | 0.02 | **0.03*** | 2244.3 | 0.76 | 0.84 | 0.46 | 1.97 | 2.48 | 0.23 |
| 4533.5 | 0.10 | 0.07 | 0.30 | 2285.3 | 0.52 | 0.62 | 0.17 | 0.69 | 0.68 | 0.86 |
| | | | | 2326.3 | 0.10 | 0.15 | 0.054 | 0.34 | 0.37 | 0.60 |
| | | | | 2360.3 | 0.06 | 0.10 | 0.37 | 0.08 | 0.12 | **0.04*** |
| | | | | 2377.3 | 0.12 | 0.16 | 0.08 | 0.14 | 0.21 | **0.048*** |
| | | | | 2390.3 | 0.09 | 0.13 | 0.38 | 0.18 | 0.35 | **0.03*** |
| | | | | 2396.3 | 2.31 | 1.83 | 0.07 | 3.21 | 3.30 | 0.83 |
| | | | | 2418.4 | 0.62 | 0.69 | 0.40 | 0.60 | 0.72 | 0.33 |
| | | | | 2431.4 | 0.07 | 0.08 | 0.59 | 0.09 | 0.29 | 0.06 |
| | | | | 2448.3 | 0.22 | 0.23 | 0.59 | 0.35 | 0.42 | 0.29 |
| | | | | 2459.4 | 3.99 | 3.76 | 0.60 | 5.39 | 3.83 | **0.04*** |
| | | | | 2489.4 | 0.06 | 0.08 | 0.12 | 0.08 | 0.08 | 0.91 |
| | | | | 2530.4 | 0.09 | 0.13 | **0.016*** | 0.16 | 0.14 | 0.40 |
| | | | | 2564.5 | - | - | - | 0.09 | 0.12 | 0.09 |
| | | | | 2592.5 | 0.30 | 0.36 | 0.23 | 0.13 | 0.17 | 0.22 |
| | | | | 2600.4 | - | - | - | 0.05 | 0.06 | 0.36 |
| | | | | 2605.5 | 0.09 | 0.13 | 0.41 | 0.12 | 0.34 | 0.06 |
| | | | | 2622.5 | 0.15 | 0.19 | 0.12 | 0.20 | 0.26 | 0.20 |
| | | | | 2635.4 | 0.08 | 0.11 | 0.26 | 0.13 | 0.28 | 0.08 |
| | | | | 2646.5 | 0.17 | 0.24 | 0.50 | 0.38 | 0.59 | 0.06 |
| | | | | 2663.5 | 0.17 | 0.20 | 0.38 | 0.18 | 0.17 | 0.80 |
| | | | | 2704.5 | 0.31 | 0.41 | 0.08 | 0.59 | 0.51 | 0.46 |
| | | | | 2779.6 | 0.11 | 0.16 | 0.48 | 0.09 | 0.12 | 0.12 |
| | | | | 2796.6 | 0.05 | 0.09 | 0.12 | 0.06 | 0.07 | 0.34 |
| | | | | 2809.5 | - | - | - | 0.06 | 0.09 | 0.07 |
| | | | | 2837.6 | 1.18 | 1.11 | 0.66 | 0.75 | 0.73 | 0.86 |
| | | | | 2891.6 | 0.08 | 0.11 | 0.46 | 0.22 | 0.34 | 0.12 |
| | | | | 2908.6 | 0.06 | 0.08 | 0.11 | 0.11 | 0.08 | 0.51 |
| | | | | 2925.6 | - | - | - | 0.04 | 0.03 | 0.68 |
| | | | | 2966.6 | 0.05 | 0.09 | 0.45 | 0.08 | 0.16 | **0.04*** |
| | | | | 2983.6 | - | - | - | 0.03 | 0.03 | 0.79 |

| | | | | | | |
|---|---|---|---|---|---|---|
| 2996.6 | - | - | - | 0.03 | 0.06 | 0.05 |
| 3024.7 | 0.14 | 0.19 | 0.51 | 0.13 | 0.20 | 0.08 |
| 3041.7 | 0.05 | 0.07 | 0.24 | 0.06 | 0.05 | 0.81 |
| 3054.7 | - | - | - | 0.02 | 0.04 | 0.10 |
| 3082.7 | 0.15 | 0.23 | 0.16 | 0.27 | 0.27 | 0.95 |
| 3095.6 | - | - | - | 0.04 | 0.06 | 0.22 |
| 3140.8 | 0.04 | 0.06 | 0.35 | 0.03 | 0.04 | 0.22 |
| 3215.8 | 0.39 | 0.50 | 0.25 | 0.12 | 0.14 | 0.65 |
| 3228.9 | - | - | - | 0.03 | 0.03 | 0.74 |
| 3269.9 | 0.10 | 0.16 | 0.40 | 0.19 | 0.28 | 0.12 |
| 3286.8 | 0.03 | 0.05 | 0.15 | 0.06 | 0.04 | 0.60 |
| 3328.0 | 0.03 | 0.05 | 0.33 | 0.05 | 0.07 | 0.14 |
| 3402.9 | 0.15 | 0.25 | 0.37 | 0.08 | 0.10 | 0.35 |
| 3456.9 | - | - | - | 0.04 | 0.09 | 0.11 |
| 3460.9 | 0.06 | 0.10 | 0.30 | 0.10 | 0.15 | 0.11 |
| 3473.9 | - | - | - | 0.03 | 0.03 | 0.79 |
| 3514.8 | - | - | - | 0.02 | 0.02 | 0.38 |
| 3590.0 | 0.03 | 0.06 | 0.36 | 0.04 | 0.06 | 0.08 |
| 3631.3 | - | - | - | 0.02 | 0.04 | 0.14 |
| 3648.0 | 0.07 | 0.10 | 0.54 | 0.08 | 0.11 | 0.20 |
| 3764.2 | - | - | - | 0.02 | 0.04 | **0.04*** |
| 3777.1 | - | - | - | 0.02 | 0.05 | 0.10 |
| 3835.3 | 0.02 | 0.06 | 0.31 | 0.03 | 0.06 | 0.10 |
| 3852.1 | - | - | - | 0.03 | 0.03 | 0.84 |
| 3893.3 | - | - | - | 0.02 | 0.04 | 0.30 |
| 4009.0 | - | - | - | 0.02 | 0.04 | 0.09 |
| 4026.3 | 0.06 | 0.09 | 0.48 | 0.11 | 0.17 | 0.15 |
| 4039.3 | - | - | - | 0.02 | 0.02 | 0.43 |
| 4080.5 | - | - | - | 0.01 | 0.03 | 0.29 |
| 4213.4 | 0.03 | 0.04 | 0.42 | 0.06 | 0.12 | 0.05 |
| 4400.5 | - | - | - | 0.03 | 0.08 | 0.12 |
| 4574.8 | - | - | - | 0.01 | 0.01 | 0.20 |
| 4587.9 | - | - | - | 0.01 | 0.02 | 0.24 |
| 4649.6 | - | - | - | 0.01 | 0.02 | 0.26 |

**Table S4. Sex comparison of protein N- glycan abundance in plasma, cortex, and cerebellum.** For plasma samples male=8, female=6. For brain samples male=6, female=4. p-values <0.05 shown in bold. Related to Fig. 4.

**The restricted nature of protein glycosylation in the mammalian brain**

**Mealer, et al,. 2020.**

| Glycan Name | *m/z* | GlcNAc | Mannose | Galactose | NeuGc | Fucose | High-Man | Hybrid | Antenna |
|---|---|---|---|---|---|---|---|---|---|
| Man-5 | 1579.9 | 2 | 5 | 0 | 0 | 0 | 1 | 0 | 0 |
| A1G1 | 1620.9 | 3 | 3 | 1 | 0 | 0 | 0 | 0 | 1 |
| A2 | 1661.9 | 4 | 3 | 0 | 0 | 0 | 0 | 0 | 2 |
| Man-6 | 1784.0 | 2 | 6 | 0 | 0 | 0 | 1 | 0 | 0 |
| FA2 | 1836.0 | 4 | 3 | 0 | 0 | 1 | 0 | 0 | 2 |
| A2B | 1907.1 | 5 | 3 | 0 | 0 | 0 | 0 | 0 | 3 |
| Man-7 | 1988.1 | 2 | 7 | 0 | 0 | 0 | 1 | 0 | 0 |
| A1G1S1 | 2012.2 | 3 | 3 | 1 | 1 | 0 | 0 | 0 | 1 |
| FA2G1 | 2040.2 | 4 | 3 | 1 | 0 | 1 | 0 | 0 | 2 |
| Man-8 | 2192.2 | 2 | 8 | 0 | 0 | 0 | 1 | 0 | 0 |
| A1G1S1H | 2216.2 | 3 | 4 | 1 | 1 | 0 | 0 | 1 | 1 |
| A2G1S1 | 2257.3 | 4 | 3 | 1 | 1 | 0 | 0 | 0 | 2 |
| Man-9 | 2396.3 | 2 | 9 | 0 | 0 | 0 | 1 | 0 | 0 |
| A1G1S1H | 2420.3 | 3 | 5 | 1 | 1 | 0 | 0 | 1 | 1 |
| FA2G1S1 | 2431.4 | 4 | 3 | 1 | 1 | 1 | 0 | 0 | 2 |
| A2G2S1 | 2461.4 | 4 | 3 | 2 | 1 | 0 | 0 | 0 | 2 |
| FA2G2S1 | 2635.5 | 4 | 3 | 2 | 1 | 1 | 0 | 0 | 2 |
| A2G2S2 | 2852.6 | 4 | 3 | 2 | 2 | 0 | 0 | 0 | 2 |
| A3G3S1 | 2910.6 | 5 | 3 | 3 | 1 | 0 | 0 | 0 | 3 |
| FA2G2A2 | 3026.7 | 4 | 3 | 2 | 2 | 1 | 0 | 0 | 2 |
| A2G2S3 | 3243.8 | 4 | 3 | 2 | 3 | 0 | 0 | 0 | 2 |
| A3G3S2 | 3301.8 | 5 | 3 | 3 | 2 | 0 | 0 | 0 | 3 |
| FA2G2S3 | 3417.9 | 4 | 3 | 2 | 3 | 1 | 0 | 0 | 2 |
| A3G3S3 | 3693.0 | 5 | 3 | 3 | 3 | 0 | 0 | 0 | 3 |
| A3FG3S3 | 3867.1 | 5 | 3 | 3 | 3 | 1 | 0 | 0 | 3 |
| A3G3S4 | 4084.3 | 5 | 3 | 3 | 4 | 0 | 0 | 0 | 3 |
| A3FG3S4F | 4258.4 | 5 | 3 | 3 | 4 | 1 | 0 | 0 | 3 |
| A3G3S5 | 4475.3 | 5 | 3 | 3 | 5 | 0 | 0 | 0 | 3 |
| A4G4S4 | 4533.5 | 6 | 3 | 4 | 4 | 0 | 0 | 0 | 4 |

**The restricted nature of protein glycosylation in the mammalian brain**

**Mealer, et al,. 2020.**

**Table S5. Plasma protein N-glycan structure, name, mass, and characteristics.** Related to Fig. 4.

| | Plasma | | | Cortex | | | Cerebellum | | |
|---|---|---|---|---|---|---|---|---|---|
| **Glycan Category** | **Male** | **Female** | ***p-value*** | **Male** | **Female** | ***p-value*** | **Male** | **Female** | ***p-value*** |
| Paucimannose | - | - | - | 3.10 | 3.33 | 0.43 | 1.17 | 2.87 | 0.12 |
| High-mannose | 1.1 | 1.1 | 0.87 | 62.8 | 59.8 | 0.34 | 61.3 | 59.9 | 0.67 |
| Mono-antennary | 1.2 | 0.9 | 0.47 | 18.9 | 20.3 | 0.24 | 12.9 | 16.5 | 0.09 |
| Bi-antennary | 75.4 | 80.8 | **0.046*** | 13.5 | 14.2 | 0.71 | 21.7 | 17.2 | **0.047*** |
| Tri-antennary | 22.2 | 17.2 | 0.07 | 1.5 | 2.2 | 0.21 | 2.6 | 3.0 | 0.48 |
| Tetra-antennary | 0.1 | 0.1 | 0.30 | 0.09 | 0.14 | 0.46 | 0.3 | 0.6 | 0.13 |
| Hybrid | 0.8 | 0.6 | 0.46 | 5.2 | 6.3 | 0.22 | 8.3 | 12.1 | 0.07 |
| Bisected | - | - | - | 30.2 | 31.9 | 0.53 | 34.4 | 32.0 | 0.50 |
| Fucose | 9.4 | 47.7 | **0.00006***** | 34.5 | 36.9 | 0.39 | 35.2 | 34.3 | 0.72 |
| Galactose | 98.2 | 98.2 | 0.99 | 13.1 | 14.4 | 0.52 | 14.0 | 14.9 | 0.65 |
| NeuAc | - | - | - | 1.5 | 2.2 | 0.42 | 2.8 | 4.8 | **0.040*** |
| NeuGc | 97.7 | 97.4 | 0.74 | - | - | - | - | - | - |

**Table S6. Sex differences in classes of N-glycans.**
Subgroup analysis of N-glycans between sexes revealed significantly increased fucosylation and reduced branching in female plasma, as well as a minor increase in sialylation and branching in the female cerebellum. The cortex trends similarly to the cerebellum, though the differences do not reach statistical significance. For plasma samples male=8, female=6. For brain samples male=6, female=4. p-values <0.05 shown in bold. Related to Fig. 4.

**The restricted nature of protein glycosylation in the mammalian brain**

**Mealer, et al,. 2020.**

| *m/z* | Male | Female |
|---|---|---|
| 534.30 | 0.58 | 1.26 |
| 691.42 | 0.25 | 1.22 |
| 779.47 | 1.28 | 3.48 |
| 895.49 | 3.74 | 5.74 |
| 912.51 | 1.05 | 1.51 |
| 925.53 | 0.13 | 0.19 |
| 983.56 | 0.39 | 0.36 |
| 1069.62 | 0.24 | 1.26 |
| 1099.60 | 5.70 | 7.94 |
| 1129.63 | 0.22 | 0.34 |
| 1157.65 | 5.95 | 4.35 |
| 1256.69 | 68.92 | 63.22 |
| 1286.71 | 0.69 | 0.42 |
| 1316.74 | 0.18 | 0.20 |
| 1344.76 | 3.28 | 3.01 |
| 1361.79 | 0.44 | 0.36 |
| 1460.80 | 1.29 | 1.15 |
| 1518.84 | 1.33 | 0.72 |
| 1535.86 | 0.53 | 0.36 |
| 1548.85 | 0.55 | 0.44 |
| 1589.90 | 0.08 | 0.11 |
| 1617.88 | 1.50 | 1.15 |
| 1705.95 | 0.77 | 0.53 |
| 1722.96 | 0.33 | 0.23 |
| 1910.06 | 0.49 | 0.38 |
| 1978.26 | 0.10 | 0.06 |

**Table S7. Sex comparison of protein O-glycan abundance in cortex.** Male=3, Female=2. Related to Figure 4.
