## Supplementary figures and images for "The restricted nature of protein glycosylation in the mammalian brain"

### Supplemental File 1

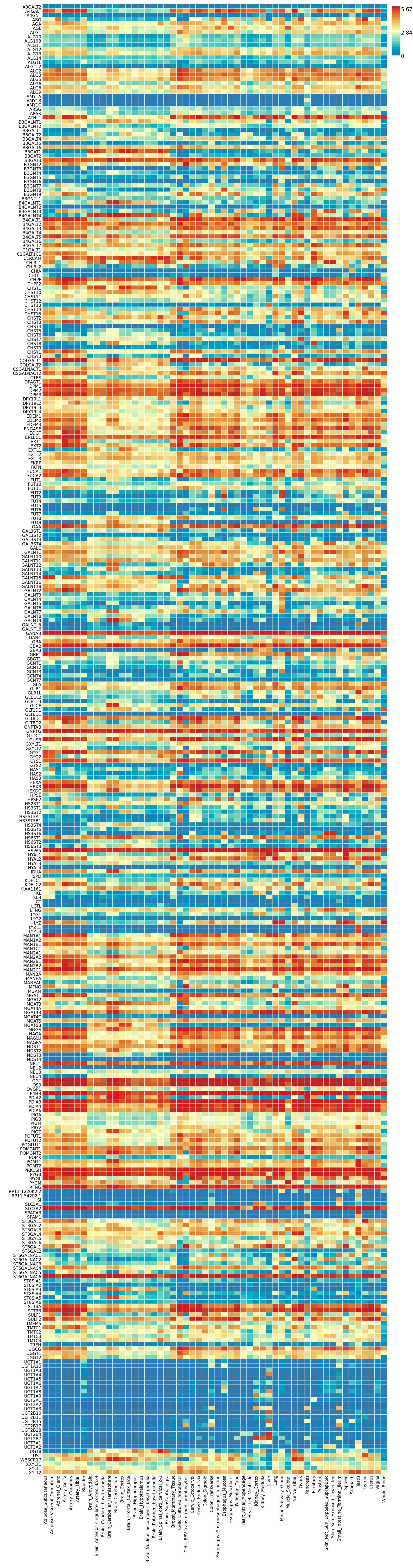
